# Protein Kinase C δ: a critical hub regulating macrophage immunomodulatory functions during *Mycobacterium tuberculosis* infection

**DOI:** 10.1101/2025.05.19.653976

**Authors:** Rudranil Hazra, Mumin Ozturk, Nashied Peton, Tariq Ganief, Sibongiseni KL Poswayo, Robert P. Rousseau, Saiyukthi Naidoo, Shelby-Sara Jones, Anca F. Savulescu, Raymond M. Moseki, Abhimanyu, Nelita Du Plessis, Jonathan Blackburn, Musa M. Mhlanga, C Ronald Kahn, Frank Brombacher, Robert J. Wilkinson, Suraj P. Parihar

**Affiliations:** Wellcome Discovery Research Platforms in Infections, Centre for Infectious Diseases Research in Africa (CIDRI-Africa), Institute of Infectious Diseases and Molecular Medicine (IDM), Faculty of Health Sciences, University of Cape Town, Cape Town 7925, Republic of South Africa; Division of Medical Microbiology, Institute of Infectious Diseases and Molecular Medicine (IDM), Department of Pathology, Faculty of Health Sciences, University of Cape Town, Cape Town 7925, Republic of South Africa; Epigenomics & Single Cell Biophysics Group, Department of Cell Biology FNWI, Radboud University, Nijmegen, Netherlands; Department of Internal Medicine, Radboud University Medical Centre, Nijmegen, The Netherlands; Drug Discovery and Development Centre (H3D), University of Cape Town, Cape Town 7925, Republic of South Africa; Division of Chemical, Systems & Synthetic Biology, Institute of Infectious Diseases & Molecular Medicine (IDM), Faculty of Health Sciences, University of Cape Town, Cape Town, South Africa; Department of Microbiology, Biological Sciences Division, University of Chicago, Chicago, IL, USA; DSI-NRF Centre of Excellence for Biomedical Tuberculosis Research, South African Medical Research Council for Tuberculosis Research, Division of Molecular Biology and Human Genetics, Faculty of Medical and Health Sciences, Stellenbosch University, Cape Town, Republic of South Africa; Section of Integrative Physiology and Metabolism, Joslin Diabetes Center, Harvard Medical School, Boston, MA, USA; Division of Immunology, Institute of Infectious Diseases and Molecular Medicine (IDM), Department of Pathology, Faculty of Health Sciences, University of Cape Town, Cape Town 7925, Republic of South Africa; Institute of Infectious Diseases and Molecular Medicine (IDM), Department of Medicine, University of Cape Town, Cape Town, South Africa; The Francis Crick Institute, London NW1 1AT, UK and Department of Infectious Diseases, Imperial College London, London W12 ONN, UK

## Abstract

A host-modulating candidate gene involved in putative pathogen-killing pathways, with potential novel therapeutic intervention, Protein Kinase C – δ (PKCδ) has been recognized as a critical marker of inflammation with clinical and experimental evidence in recent years. Pulmonary microenvironment during *Mtb* infection is largely governed by lung resident macrophages, initiating innate and subsequent adaptive immune responses. We investigated the role of PKCδ in macrophages using a macrophage-specific PKCδ knockout mice model (LysM^cre^PKCδ^flox/flox^). PKCδ deficiency in macrophages triggers an early lymphocytic immune response, increases neutrophil recruitment, and reduces inflammatory macrophages in the lungs, leading to higher *Mtb* burden and exacerbated pathology. Experimental and omics analysis further revealed that dysregulation of antimicrobial effector functions is detrimental to macrophage’s ability to restrict bacterial growth *in vitro*. Importantly this defect was mitigated by exogenous GM-CSF supplementation and/or overexpressing PKCδ in macrophages. Thus, PKCδ plays a crucial role in immune modulation during *Mtb* infection with GM-CSF amongst several downstream pathways through which PKCδ exerts its regulatory effects.

**Teaser:** PKCδ is crucial for immune modulation during *Mtb* infection revealing macrophages as a potential axis of signaling.

## Introduction

Tuberculosis (TB) remains a major health concern worldwide, despite significant advances in treatment and prevention strategies. Reduced access to TB diagnosis and treatment due to the COVID-19 pandemic resulted in a steep increase in TB deaths, according to the Global Tuberculosis Report 2023(*1*). However, the emergence of multi-drug resistant (MDR) strains, limited vaccine efficacy of BCG, prolonged TB treatment regimens, and socio-economic challenges persist as significant factors for failure of TB control(*2*). Genetic resistance and phenotypical tolerance of the *Mycobacterium tuberculosis* (*Mtb*) pathogen against antimycobacterial underscores an urgency to develop adjunctive anti-TB therapies to mitigate the pulmonary pathology caused by TB disease(*3, 4*). Exploiting host-pathogen interplay by an alternative approach, host-directed therapies (HDT) have been a widely studied strategy in the field of TB research(*3*). These studies showed improved efficacy for bacterial killing, strengthening innate and memory responses, disrupting the structural integrity of TB granuloma, and modulating inflammatory responses(*3, 5*). Hence, identifying potential HDT targets and understanding their involvement in key immune responses during *Mtb* infection is a foundational step toward a comprehensive understanding of the innate and adaptive components of the host defense mechanisms(*3–5*).

Protein kinase C - δ (PKCδ) is a phospholipid-dependent serine/threonine kinase first described by Nishizuka and colleagues, and it plays a critical role in intracellular signal transduction(*6–8*). As a result of its ability to phosphorylate a variety of target proteins necessary for cellular functions such as signal transduction(*9*), apoptosis(*10*), proliferation and survival(*11*), transcription(*12*), hormonal regulation(*13*), and immunological responses(*14–16*), PKCδ is considered as a key hub for immunoregulatory functions among other PKC isoforms. PKCδ null mouse (PKCδ^-/-^) exhibited B-cell accumulation(*17*) and systemic autoimmunity(*18*) with no apparent developmental and fertility issues. PKCδ is ubiquitously expressed in lymphocytes and most mammalian tissue and cell types, including myeloid cells such as DC andmacrophages(*19, 20*). However, the characterization of the PKCδ^-/-^ mouse was focused to lymphoid progenitor development in which only B cell functionality was emphasized along with the recruitment of T lymphocytes and bone marrow cells in the spleen, lymph nodes, and thymus(*18*). Previous studies using PKCδ^-/-^ mouse indicated immune protective roles of the kinase (*14–16*) against different infectious diseases. Also, high susceptibility to *Mtb* infection with increased bacterial burden, mortality, weight loss, exaggerated lung inflammation, and unrestrained cytokine response has previously been demonstrated using PKCδ^-/-^ mice model(*16*). Protective immunity to TB is thought to result from a T-cell-dependent immune response, with CD4^+^ T cells playing a vital role(*21, 22*). While PKCδ^-/-^ mice exhibit conventional T-cell-independent susceptibility in response to *Mtb* infection(*16*). Recently, experimental and clinical evidence has increasingly highlighted the importance of innate cells in protective immunity(*23, 24*). The initial uptake of *Mtb* by macrophages, the primary innate immune cell also possesses T-cell-independent, essential bactericidal activity(*25, 26*) which is also apparent in PKCδ^-/-^ mice with increased activated macrophages in the lungs during acute *Mtb* infection(*16*). However, these findings only emphasized the outcome of germ-line deficiency of PKCδ in mice which prompted us to investigate the role of macrophage-specific PKCδ deficiency during TB disease which has not hitherto been studied. We generated a macrophage-specific PKCδ knockout mouse model (LysM^cre^PKCδ^flox/flox^) by crossing lysozyme M – Cre (LysM^cre^) mice with loxP flanked PKCδ mice(*27, 28*). LysM^cre^ mice enable specific and efficient Cre-mediated deletion of lox-P flanked target genes in myelomonocytic cells including granulocytes, monocytes, and macrophages(*29*).

In this study, we demonstrated the impact of macrophage-specific PKCδ deletion on various immune cell populations across organs, lung cytokine and chemokine levels to better understand the functions as well as the immunomodulatory effect of PKCδ in TB using LysM^cre^PKCδ^flox/flox^ murine model. Our results suggest that PKCδ deficiency in macrophages exhibits an increase in mycobacterial burden in the lungs and more disseminated bacteria in the spleen at the chronic stage. Surprisingly, the lung immune cell profiling revealed T-cell-dependent host-defense through increased frequencies of T memory subsets and exhaustion markers during the initial stages of *Mtb* infection. Also, increased lung cell numbers were associated with infiltration of neutrophils at the acute stage of *Mtb* infection. In addition, we also showed that reduced *Mtb* restrictive interstitial macrophages (IM) are associated with increased bacterial load in the lungs of LysM^cre^PKCδ^flox/flox^ mice. Despite a significant reduction in granulocyte-macrophage colony-stimulating factor (GM-CSF) levels -a critical cytokine for alveolar macrophage (AM) maintenance-within the lungs of LysM^cre^PKCδ^flox/flox^ mice during *Mtb* infection, the frequencies of permissive AM remained unchanged(*30*).

To examine the macrophage-intrinsic role of PKCδ in *Mtb* infection in greater detail, we investigated macrophages from LysM^cre^PKCδ^flox/flox^ mice. Our findings mirrored the increased susceptibility of LysM^cre^PKCδ^flox/flox^ mice to *Mtb* infection in isolated macrophages. Interestingly, overexpression of PKCδ in a murine macrophage cell line, on the other hand, reduced the mycobacterial load. Further investigation revealed that PKCδ-deficient macrophages exhibited altered antimicrobial effector functions, with an elevated proinflammatory cytokine response and inflammasome activation during *Mtb* infection. Notably, we identified cellular metabolic ATP production dysregulation in the absence of PKCδ in macrophages during *Mtb* infection, which was reversed by the exogenous addition of GM-CSF, leading to reduced bacterial replication. Additionally, the transcriptomic and proteomic analysis identified that PKCδ regulates critical genes and proteins involved in immune signaling during *Mtb* infection. Specifically, its deficiency impaired GM-CSF signaling, resulting in hyperactive type I IFN signaling and neutrophil migration programs. A comprehensive macrophage signaling atlas further underscored the central role of PKCδ in modulating these pathways. Clinically, increased PKCδ expression at site of disease from active TB patients, while knockdown of PKCδ in human monocyte-derived macrophages (MDM) recapitulated the phenotype of increased bacterial replication. Overall, the inflammatory responses elicited by PKCδ demonstrated that this novel kinase is a crucial hub for orchestrating immunoregulatory functions in macrophages during *Mtb* infection.

## Results

### PKCδ deficiency in macrophages increased *Mtb* burden and worsened lung pathology *in vivo*

To generate a myeloid-specific knockout mice model, we have followed the conventional LysM^cre^-loxP genetic manipulation technique described previously by Jiayuan Shi *et al*(*31*). We confirmed the integrity of loxP-floxed (fl/fl) PKCδ in the genomic DNA extracted from the ear clips of PKCδ^flox/flox^ mice compared to wildtype(n/n) mice and the presence of Cre recombinase transgene (LysM^cre^) in LysM^cre^PKCδ^flox/flox^ (LysM^cre+^) mice compared to PKCδ^flox/flox^ (LysM^cre-^) mice, shown as linear DNA bands on the agarose gel (**Supplementary Figure 1A**). To determine the specificity and efficiency of LysM^cre^-loxP mediated cell-specific conditional knockout, we sorted different immune cell populations by flow cytometry to confirm the deletion of PKCδ specifically in macrophages while the expression of PKCδ maintained in other myeloid cell types by qRT-PCR (**Supplementary Figure 1B-C**). We also found that the PKCδ mRNA transcript was undetectable and subsequent translation of the protein was reduced in the bone-marrow-derived macrophages (BMDM) isolated from LysM^cre^PKCδ^flox/flox^ mice (**Supplementary Figure 1D-E**). Additionally, to establish a PKCδ exclusive myeloid knockout setting, we have determined the effect of PKCδ deletion on different PKC family (classical PKC, novel PKC, and atypical PKC) isoforms in BMDMs through mRNA expression by qRT-PCR (**Supplementary Figure 1F**). To identify alterations in the lung and spleen physiology of LysM^cre^PKCδ^flox/flox^ mice, we processed single-cell suspension from LysM^cre^PKCδ^flox/flox^ mice. Besides having a minor reduction in lung B cell percentage, no significant irregularities of lymphoid and myeloid immune cell frequencies were observed in the lungs or spleen of LysM^cre^PKCδ^flox/flox^ mice compared to PKCδ^flox/flox^ mice (**Supplementary Figure 1G-M**). Importantly, Cre-mediated deletion of PKCδ did not influence basal cytokine, chemokine levels, and growth factors, indicating no immunological distress in LysM^cre^PKCδ^flox/flox^ mice compared to PKCδ^flox/flox^ mice in naïve state (**Supplementary Figure 1N-Q**).

Our previous findings indicated a greater susceptibility and heightened inflammation caused by *Mtb* (H37Rv) infection in the lungs of PKCδ^-/-^ mice mediated through suboptimal macrophage functions, leading to increased pulmonary cell recruitment and enumerated *Mtb* colonies(*16*). To extend these findings and investigate clinical *Mtb* strain specific events, we intranasally challenged mice with hypervirulent *Mtb* (HN878) for 4 weeks (acute stage; 4WPI) and 12 weeks (mid-chronic stage; 12 WPI) (**Figure 1A**). We found significantly increased *Mtb* burden in the lungs of LysM^cre^PKCδ^flox/flox^ mice at both 4- and 12 WPI with dissemination to the spleen at 12 WPI (**Figure 1B-C**). An increased pulmonary immune cell infiltration was observed at 4 WPI; however, the lung weight index remained unchanged. This was accompanied by a progressively increasing *Mtb* burden, indicating sustained bacterial replication despite the increased immune response (**Supplementary Figure 2A-B**). Mycobacterial growth often correlates with the association of tissue morphology in response to infection. Therefore, we investigated whether the increased mycobacterial burden exacerbates lung inflammation in the LysM^cre^PKCδ^flox/flox^ mice by histopathological analysis. As expected, we found increased pathology defined as less free alveolar space in the lungs of LysM^cre^PKCδ^flox/flox^ mice compared to PKCδ^flox/flox^ mice at 4- and 12 WPI (**Figure 1D-E**). An inflammatory and permissive environment for mycobacterial growth was further indicated by reduced nitric oxide (NO) production, as evidenced by iNOS^+^ area in the lungs of LysM^cre^PKCδ^flox/flox^ mice at 4 WPI compared to PKCδ^flox/flox^ controls (**Figure 1F-G**). Previous studies highlighted the unique functions of GM-CSF, where it promotes host defense (*32*) but can also paradoxically compromise immune protection against *Mtb*(*33*). IL-2 plays a crucial role in shaping the T cell fate in response to *Mtb* antigens(*34*), while IL-10, is an established potent anti-inflammatory cytokine (*35*). Surprisingly, reduced levels of IL-2 at 12 WPI and GM-CSF at both 4- and 12 WPI were detected in the lung of LysM^cre^PKCδ^flox/flox^ mice compared to PKCδ^flox/flox^ mice, contributing to increased mycobacterial growth (**Figure 1H-I**). Notably, IL-10 levels were also reduced at both 4 WPI and 12 WPI in LysM^cre^PKCδ^flox/flox^ lungs. Since IL-10 is critical to limit excessive inflammation, its decreased levels could lead to dysregulated immune responses, potentially exacerbating lung inflammation(*36*). Although no significant differences in the levels of the indicated pro-inflammatory cytokines and chemokines were observed, the altered balance of IL-10, GM-CSF, and IL-2 likely contributes to the observed increase in *Mtb* burden and lung pathology (**Supplementary Figure 2C-H**).

**Figure 1.**
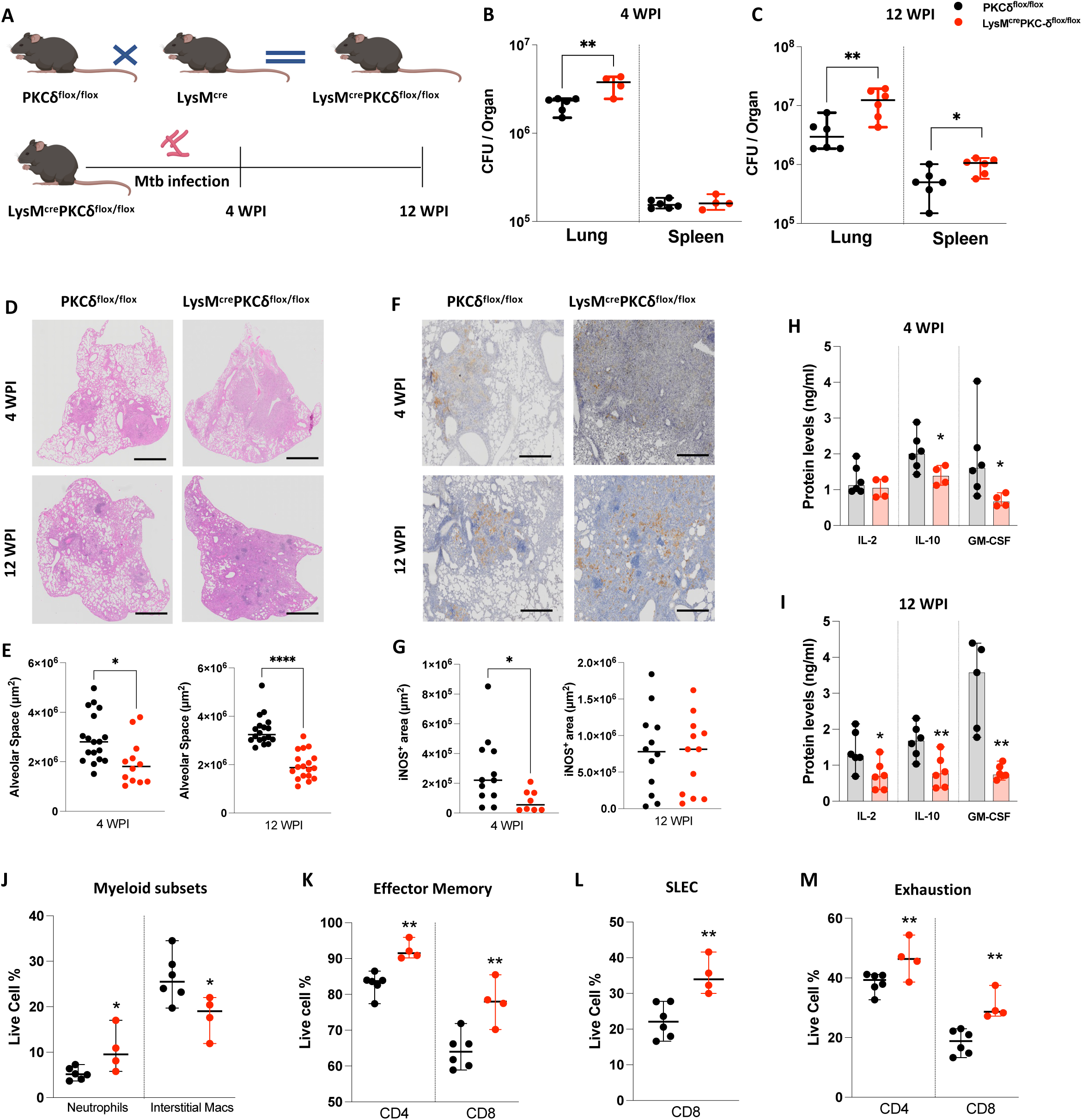
PKCδ deficiency in macrophages increased *Mtb* burden and worsened lung pathology *in vivo.* [A]. Schematic of LysM^cre^PKCδ^flox/flox^ mice generation and experimental outline of time course study. HN878 *Mtb* strain was challenged by intranasal route to LysM^cre^PKCδ^flox/flox^ and PKCδ^flox/flox^ mice at 150 CFU/mouse. **[B-C]** Animals were euthanized to determine lung/spleen CFU burden at acute stage (4 wpi) and chronic stage (12 wpi) of Mtb infection. **[D-E]** Quantification of alveolar space represented with their respective (40X) scanned images (Scale bar = 1000 µm). **[F-G]** Quantification of iNOS+ area represented with their respective (40X) scanned images (Scale bar = 100 µm). **[H-I]** Determination of IL-2, IL-10 and GM-CSF levels by ELISA in the lung homogenates of LysM^cre^PKCδ^flox/flox^ and PKCδ^flox/flox^ mice at 4 wpi and 12 wpi *Mtb* infection. **[J-M]** Flow cytometry analysis of myeloid subsets, CD4/CD8 effector memory (CD44+CD62L-), CD8 short lived effector cells (SLEC) (KLRG1+CD127-), and CD4/CD8 Exhaustion (PD1-KLRG1+) reported as frequency of live cells at 4 wpi of *Mtb* infection. All data shown mean+/-SD and is representative of two independent experiments with n=4-6 mice/group. Statistical analyses were performed using an unpaired student t-test. Asterisks are defining significance compared to the control group as: *p < 0.05, **p < 0.01, ****p < 0.0001.

The complexity of myeloid cell subsets that control TB-induced inflammation via distinct unique immune responses may result in protective or provide a favorable environment for pathogen survival(*37*). Susceptibility of PKCδ^-/-^ mice to *Mtb* infection has previously been associated with increased involvement of activated macrophages along with concomitant reduction in alveolar macrophage and DC subsets(*16*). Based on the mycobacterial burden and increased pulmonary cell numbers during acute *Mtb* infection in the LysM^cre^PKCδ^flox/flox^ mice, we further investigated the cell frequencies of various myeloid cells in the absence of PKCδ in macrophages. Our findings revealed an increased influx of neutrophils early at 4 WPI in the LysM^cre^PKCδ^flox/flox^ mice when compared to PKCδ^flox/flox^ mice (**Figure 1J**). This indicates an increased pro-inflammatory response given existing evidence of functional versatility and phenotypic heterogeneity of neutrophils with regard to PKCδ inhibition(*38*). We did not observe any differences in the frequencies of pulmonary eosinophils, monocytes, and DC subsets in the LysM^cre^PKCδ^flox/flox^ mice compared to PKCδ^flox/flox^ mice (**Supplementary Figure 2I-J**). However; ontologically distinct macrophage subsets exhibited differential effects in the absence of PKCδ with a significant reduction in the frequencies of interstitial macrophages (IM) (**Figure 1J**), which are associated with pathogen restriction(*39*). In contrast, alveolar macrophages (AM), known to be permissive to the pathogen, remained intact at 4 WPI of *Mtb* infection (**Supplementary Figure 2K**). Notably, a gradual increase of pulmonary IM was observed in the LysM^cre^PKCδ^flox/flox^ mice at 12 WPI (**Supplementary Figure 2L**). While the early encounter of *Mtb* with macrophages are a naturally occurring host defence mechanism upon infection, these cells play a pivotal role in triggering T cell responses(*40*). Previously, PKCδ^-/-^ mice showed no variability in conventional T cell recruitment during *Mtb* infection(*16*), however, T memory cell responses were not investigated. Flow cytometry analysis revealed that CD4^+^ and CD8^+^ effector memory and CD8^+^ short-lived effector cell (SLEC) frequencies were significantly increased in LysM^cre^PKCδ^flox/flox^ mice at 4 WPI (**Figure 1K-L**). In murine TB, KLRG1 but not PD-1 is recognized as a superior marker for T cell differentiation to exhaustion(*41*). The observed aberrations in memory and effector T cell subsets were most likely driven by the exhaustion of CD4^+^/CD8^+^ T cells characterized by increased PD-1^-^KLRG1^+^ frequencies, in response to high antigen loads in LysM^cre^PKCδ^flox/flox^ mice at earlier stages of infection (**Figure 1M**). However, no marked increase was observed in the frequencies of lymphocytes including conventional T cells except for Natural Killer (NK) cells at 4 WPI in LysM^cre^PKCδ^flox/flox^ mice, when compared to PKCδ^flox/flox^ mice including at later stages of progressive *Mtb* infection (**Supplementary Figure 2M-S**). Collectively, these results showed that the deletion of PKCδ in macrophages worsened disease outcome with exaggerated lung pathology, early lymphocytic immune exhaustion, and dysregulated myeloid subsets, ultimately promoting increased bacterial burden and dissemination *in vivo*.

### Ablation of PKCδ in macrophages affects varying antimicrobial effector functions *in vitro*

Based on the above role of macrophage-specific PKCδ mediated immune modulation during *Mtb* infection *in vivo*, we analysed whether macrophages from LysM^cre^PKCδ^flox/flox^ mice exerted similar effects *in vitro*. As expected, we found a significant increase in mycobacterial growth at 48 hour-post infections (hpi) in BMDM of LysM^cre^PKCδ^flox/flox^ compared to PKCδ^flox/flox^ (**Figure 2A**). To further extend our insights into the role of macrophage-specific PKCδ in sustaining immune responses during *Mtb* infection, we overexpressed PKCδ in RAW264.7 murine macrophage cell line to assess whether it could rescue the phenotype of PKCδ deficiency (**Supplementary Figure 3A**). Indeed, we found a significant decrease in mycobacterial growth in PKCδ overexpressed cells compared to the control cells at 72 and 144 hpi (**Figure 2B**). These results show that PKCδ critically regulates macrophage-mediated protective functions during *Mtb* infection. To reduce *Mtb* survival, macrophages use reactive oxygen species (ROS) to restrict the growth of cytosolic bacteria which escape from the phagolysosomal compartment, a primary component of their antibacterial defense (*42, 43*). Despite increased bacterial growth, surprisingly, we found elevated levels of cellular and mitochondrial ROS production at 24 and 48 hpi in BMDM of LysM^cre^PKCδ^flox/flox^ compared to PKCδ^flox/flox^ (**Figure 2C-D**). Also, previous research has shown the consecutive fusion of *Mtb* trapped in phagosome with the lysosome is required for microbicidal activity, leading to bacterial destruction inside macrophages(*33*). We determined that phagosome acidification was slightly reduced at 4 hpi in LysM^cre^PKCδ^flox/flox^ compared to PKCδ^flox/flox^ macrophages, however, no major effect overall on acidification of phagosomes using pHrodo was observed (**Supplementary Figure 3B**). Inducible nitric oxide synthase (iNOS), a major nitric oxide (NO) producing enzyme, is critical for the macrophage antimicrobial activity(*44*). qRT-PCR results revealed that iNOS expression in LysM^cre^PKCδ^flox/flox^ was reduced at 24 and 48 hpi with subsequently reduced levels of NO production at 48 hpi compared to PKCδ^flox/flox^ BMDM (**Figure 2E-F**).

**Figure 2.**
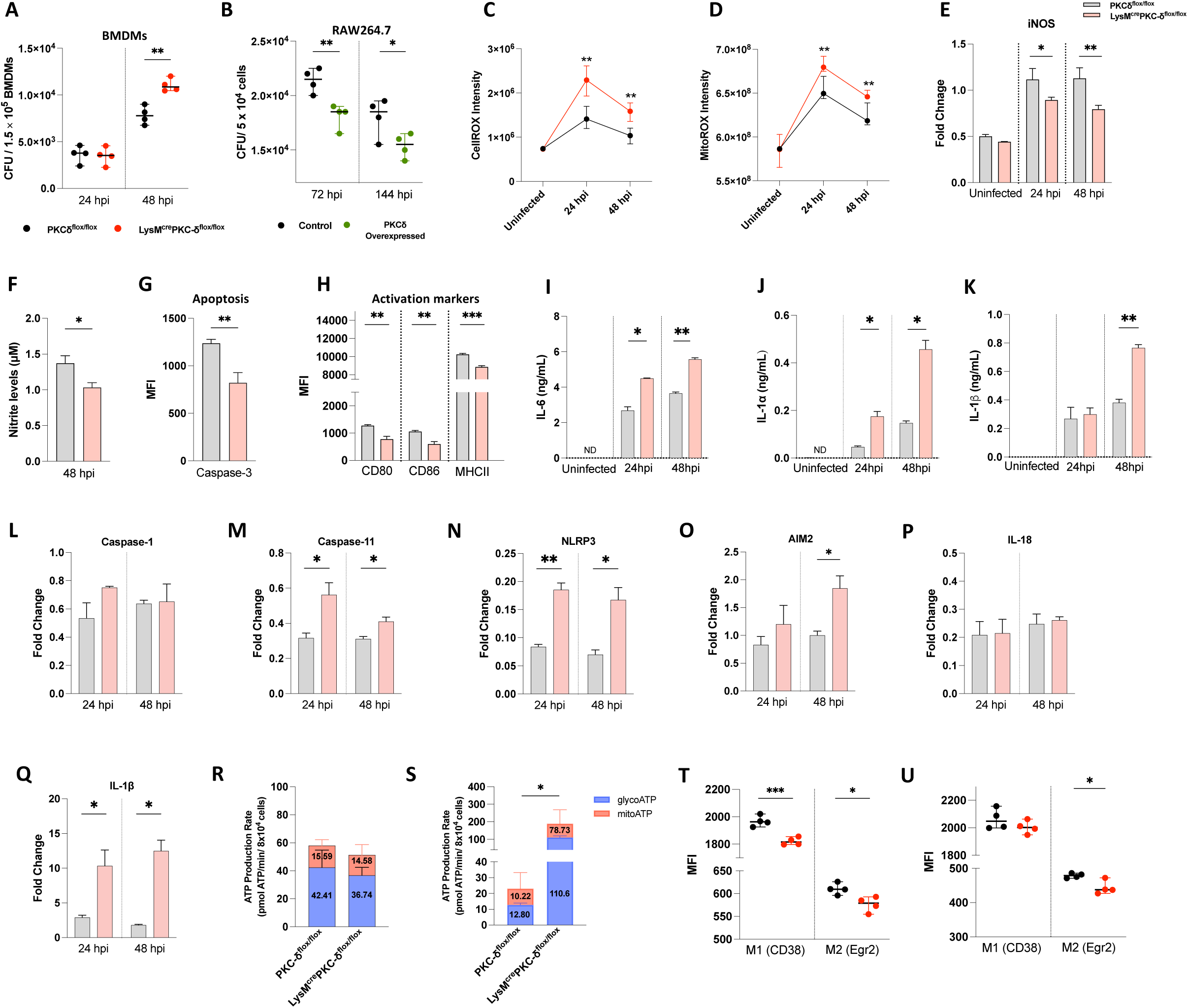
Ablation of PKCδ in macrophages led to varying antimicrobial effector functions *in vitro.* [A]. Determination of mycobacterial burden in bone-marrow-derived macrophages from LysM^cre^PKCδ^flox/flox^ and PKCδ^flox/flox^ mice at indicated time points. **[B]** Determination of mycobacterial burden in PKCδ-overexpressed RAW264.7 murine macrophage cells at indicated time points. **[C-D]** Accumulation of cellular and mitochondrial ROS were determined at indicated time points in bone-marrow-derived macrophages from LysM^cre^PKCδ^flox/flox^ and PKCδ^flox/flox^ mice during *Mtb* infection. **[E]** iNOS expression and subsequent **[F]** nitrite levels in bone-marrow-derived macrophages from LysM^cre^PKCδ^flox/flox^ and PKCδ^flox/flox^ mice at indicated time points. **[G-H]** Determination of Mean Fluorescence Intensity (MFI) of apoptotic marker Caspase-3 and macrophage activation markers (CD80, CD86, MHCII) by flow cytometry in bone-marrow-derived macrophages from LysM^cre^PKCδ^flox/flox^ and PKCδ^flox/flox^ mice at 24 hours post *Mtb* infection. **[I-K]** Determination of pro-inflammatory cytokine (IL-1α, IL-1β, IL-6) levels in bone-marrow-derived macrophages from LysM^cre^PKCδ^flox/flox^ and PKCδ^flox/flox^ mice in *Mtb* infection. **[L-Q]** mRNA expression of inflammasome pathway mediators (Caspase-1, Caspase-11, NLRP3, AIM2, IL-18 and IL-1β) in *Mtb*-infected bone-marrow-derived macrophages from LysM^cre^PKCδ^flox/flox^ and PKCδ^flox/flox^ mice. **[R-S]** Total ATP production rate and metabolic dependence of bone-marrow-derived macrophages from LysM^cre^PKCδ^flox/flox^ and PKCδ^flox/flox^ mice at naive state and 24 hours post *Mtb* infection. **[T-U]** Determination of Mean Fluorescence Intensity (MFI) of macrophage phenotypic M1 and M2 marker by flow cytometry in bone-marrow-derived macrophages from LysM^cre^PKCδ^flox/flox^ and PKCδ^flox/flox^ mice at 24 hours post *Mtb* infection. All data shown mean+/- SD and is representative of two independent experiments performed in quadruplicate. Statistical analyses were performed using an unpaired student t-test. Asterisks are defining significance compared to the control group as: *p < 0.05, **p < 0.01, ***p < 0.001.

Previously, it has been shown that siRNA inhibition of PKCδ in vascular smooth muscle cells (SMC) restricts both initiation and execution of caspase-3-mediated apoptosis(*45*). A similar reduction in caspase-3 activity was observed in LysM^cre^PKCδ^flox/flox^ BMDM at 24 hpi (**Figure 2G**). In addition, we found that bacterial proliferation was associated with impaired macrophage activation via reduced surface expression of activation markers CD80, CD86, and major histocompatibility complex II (MHCII) in LysM^cre^PKCδ^flox/flox^ BMDM (**Figure 2H**). Unlike in the *in vivo* settings, we found a significant increase in IL-6, IL-1α, and IL-1β production in the BMDM of LysM^cre^PKCδ^flox/flox^ compared to PKCδ^flox/flox^ mice at the indicated time points during *Mtb* infection (**Figure 2I-K**). The release of pro-inflammatory cytokines IL-1β and IL-18 are governed by either canonical or non-canonical inflammasomes, which serve as important signalling platforms to detect pathogenic bacteria (*46, 47*). PKCδ-mediated inflammasome activation in macrophages appeared primarily dependent on the caspase-11-NLRP3 non-canonical axis, with AIM2-mediated activation and not on caspase-1 (**Figure 2L-O**). As expected, we also found higher expression of IL-1β but not IL-18 in LysM^cre^PKCδ^flox/flox^ compared to PKCδ^flox/flox^ macrophages during *Mtb* infection (**Figure 2P-Q**). The pathogenic success of *Mtb* is also connected to its capacity to recalibrate metabolic processes in infected macrophages (*48*). To assess the energy metabolism, we measured the adenosine triphosphate (ATP) synthesis rate by LysM^cre^PKCδ^flox/flox^ BMDMs, as ATP is necessary for P2X purinergic receptor 7 (P2X7) mediated activation of the inflammasome and subsequent IL-1β production (*49–51*). Utilizing the Seahorse Bioanalyzer, we found that the total ATP production rate was unchanged in naïve macrophages but drastically increased in LysM^cre^PKCδ^flox/flox^ BMDM during *Mtb* infection (**Figure 2R-S**). Additionally, we found that the enhanced ATP production in LysM^cre^PKCδ^flox/flox^ BMDM during *Mtb* infection induced a relative increase in glycolysis than oxidative phosphorylation (OXPHOS)/ mitochondrial origin of cellular energy metabolism. Our previously published Cap Analysis Gene Expression (CAGE)-transcriptomics dataset (*52*) demonstrated predominant PKCδ expression in M2 (IL-4 stimulated) macrophages, a finding we further confirmed by qRT-PCR in IL-4 stimulated BMDM during *Mtb* infection (**Supplementary Figure 3C)**. This is in agreement with a recent study reporting PKCδ expression in M2-like BMDM in cancer(*53*). However, how the absence of PKCδ influences macrophage polarization has not been explored. Using two novel and improved markers (CD38 & Egr2)(*54*) to distinguish distinct murine macrophage phenotypes, we found that BMDM from LysM^cre^PKCδ^flox/flox^ mice displayed significantly reduced surface expressions of both M1 and M2 markers compared to PKCδ^flox/flox^ BMDM at naive state (**Figure 2T**). During *Mtb* infection, the surface expression of M1 marker was restored, whereas M2 marker expression remained reduced (**Figure 2U**), further supporting the association of PKCδ with M2-like macrophages. These findings collectively suggest that PKCδ in macrophages orchestrates a variety of antimicrobial effector activities, activation and polarization during *Mtb* infection.

### Loss of PKCδ in macrophages alters IFN signalling and chemotaxis at the transcriptomic level

To investigate pathways underlying the pathogenesis during *Mtb* infection in the absence of PKCδ in macrophages, we performed bulk RNA-Seq to examine differentially expressed genes (DEG) in BMDM from LysM^cre^PKCδ^flox/flox^ mice and compared them with PKCδ^flox/flox^ BMDM at 0 hr uninfected and 24 hr post *Mtb* infection (**Figure 3A**). We identified a total of 100 significant DEG at 0 hr uninfected timepoint, among which 26 genes were downregulated and 74 genes were upregulated in LysM^cre^PKCδ^flox/flox^ BMDM (**Figure 3B**). We observed a modest transcriptional response with 53 significant DEG at 24 hr post *Mtb* infection, among which 38 genes were downregulated and 15 genes were upregulated in LysM^cre^PKCδ^flox/flox^ BMDM (**Figure 3C**). Several of these DEG aligned with our *in vivo* and *in vitro* findings, supporting the observed regulatory effects. As expected, both at 0 hr uninfected and 24 hr post *Mtb* infection, significant downregulation of Lyz2 was identified in LysM^cre^PKCδ^flox/flox^ BMDMs (**Figure 3B-C**). This indicates that endogenous reduction of Lysozyme (Lyz2) gene function as the consequence of floxed-PKCδ deletion in macrophages by Cre recombinase was sustained throughout our experiments. In agreement with our previous findings, we also observed a significant reduction of Nos2 expression in LysM^cre^PKCδ^flox/flox^ BMDM during *Mtb* infection (**Figure 3C**). Additionally, *Mtb* infection led to broad downregulation of multiple interferon-stimulated genes (ISG) in LysM^cre^PKCδ^flox/flox^ BMDM, including but not limited to Ifi208, Cmpk2, Ifit2, Gbp5, Fgl2. Studies have also shown that Lipocalin-2 (Lcn2), a regulator of iron sequestered bacterial growth, promotes neutrophil recruitment by inducing granulocyte colony-stimulating factor (G-CSF) levels by alveolar macrophages during *Mtb* infection(*55*). We found both Lcn2 and G-CSF (CSF3) were significantly increased in LysM^cre^PKCδ^flox/flox^ BMDMs during *Mtb* infection (**Figure 3C**). This increase suggests enhanced expression of neutrophil recruitment-associated genes in macrophages, potentially linking the observed *in vitro* transcriptional changes to the *in vivo* phenotype. Moreover, Gene Ontology (GO) analysis revealed several DEG enriched pathways related to the proliferation of leukocytes and mononuclear cells at naive state (**Supplementary Figure 4A**), whereas pathways significantly over-represented at 24 hr post *Mtb* infection predominantly included neutrophil migration (**Figure 3D**). We also observed a subset of genes associated with neutrophil migration were significantly altered in LysM^cre^PKCδ^flox/flox^ compared to PKCδ^flox/flox^ macrophages at 24 hr post *Mtb* infection (**Figure 3E**). Neutrophilic inflammation contributes to tuberculosis pathology, with some subsets creating a permissive state for *Mtb* growth(*56, 57*). Additionally, nitric oxide, the main product of the Nos2 enzyme, is shown to inhibit neutrophil recruitment, which otherwise creates a nutrient-rich environment for *Mtb* growth(*58*). Previous studies found that depletion of an epigenetic reader speckled protein 140 (Sp140) in mice and specifically in macrophages exhibits a hyper type I IFN response and increased neutrophil recruitment in *Mtb* infection(*59*). Notably, we found Sp140 was significantly decreased in BMDM of LysM^cre^PKCδ^flox/flox^ compared to PKCδ^flox/flox^ during *Mtb* infection (**Figure 3C**). To examine this further, we performed immune response enrichment analysis (IREA) to infer cytokine responses based on the regulated transcriptional signature in the absence of PKCδ in macrophages(*60*). Surprisingly, IREA indicated type I and II IFNs (IFNα1, IFNβ, IFNγ) response in LysM^cre^PKCδ^flox/flox^ BMDM at 0 hr (**Supplementary Figure 4B**) and 24 hr post *Mtb* infection (**Figure 3F),** suggesting a possible fundamental role of PKCδ in macrophage triggering of IFN signaling as reported recently by Chaib and colleagues(*53*). Studies have shown that elevated Sp140 levels augment STAT1 inhibition(*61*), whereas, Sp140 blockade enriches IRF1(*62*), conducive to reducing macrophage inflammatory status. Moreover, PKCδ is known to be an upstream mediator of STAT1 phosphorylation inducing type I IFN biological response in both B and T lymphoblasts(*63*). In parallel, ChIP-X Enrichment Analysis (ChEA) analysis(*64*) revealed that both STAT1 and IRF1 were the top consensus transcription factors among others in the absence of PKCδ in macrophages at naive and in *Mtb* infection (**Figure 3H, Supplementary Figure 4C**), suggesting a possible role of other PKC isoforms regulating STAT1 and IRF1 in macrophages, as also noted by Pilz *et al*(*65*), which requires further mechanistic validation. Findings herein indicate that PKCδ is critical in regulating hyper type I IFN signaling and neutrophilic migration at the transcriptome level.

**Figure 3.**
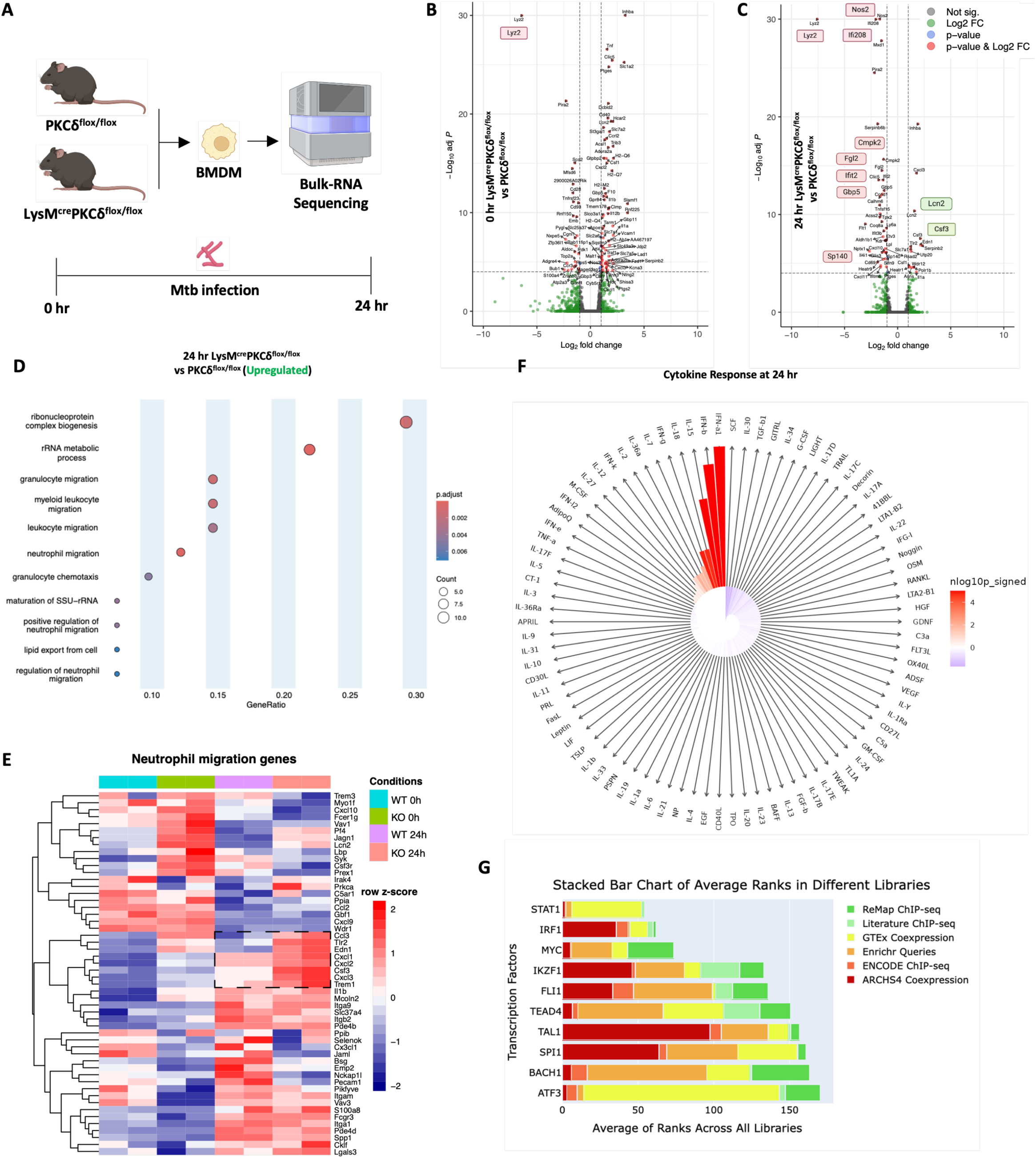
Intracellular loss of PKCδ in macrophages indicated IFN signaling and neutrophil migration at transcriptome level during *Mtb* infection. [A]. Experimental outline of bulk-RNA sequencing of BMDMs from LysM^cre^PKCδ^flox/flox^ and PKCδ^flox/flox^ mice at indicated time points. **[B-C]** Volcano plots for uninfected (0 hr) and infected (24 hr) DEGs are shown between LysM^cre^PKCδ^flox/flox^ and PKCδ^flox/flox^ BMDMs. Upregulated and downregulated genes are defined as positive p-value and Log2 FC and negative p-value and Log2 FC, respectively. Dotted line denotes threshold (Fold change ≥ 0.5; p-value ≥ 0.05) on X- and Y-axis. **[D]** Horizontal dot plots (Ora) detailing the association of enriched Gene Ontology (GO) biological processes are shown between LysM^cre^PKCδ^flox/flox^ and PKCδ^flox/flox^ BMDMs at 24 hr post *Mtb* infection. **[E]** Heatmap of DEGs cluster associated with neutrophil migration. Significant DEGs are demarcated with doted square. **[F]** IREA cytokine enrichment plot showing the enrichment score (ES) for each of the cytokine response in LysM^cre^PKCδ^flox/flox^ BMDMs at 24 hr post *Mtb* infection. Bar length is representing the ES with darker red (enriched in LysM^cre^PKCδ^flox/flox^ BMDMs) and darker blue (enriched in PKCδ^flox/flox^ BMDMs). **[G]** Horizontal bar chart representing the top ranked transcription factors at 24 hr post *Mtb* infection according to their average integrated scores across all the libraries. All data shown are analysed and produced using R studio packages and appyters web-based software. Schematic cartoons by Biorender.com.

### Dysregulated protein clusters contribute to neutrophil activation in PKCδ-deficient macrophages during *Mtb* infection

To complement investigation of the transcriptome, we extended our systemic approach by employing proteomic analysis to explore the broad spectrum of protein dysregulation and phospho-proteomics to specifically investigate the phosphorylation-dependent signalling pathways likely influenced by PKCδ deletion in macrophages at 0 min (Uninfected) and 30 min post *Mtb* infection (Infected) (**Figure 4A**). Combined heat map cluster analysis of the total proteome shows significant differences in the protein expression patterns between LysM^cre^PKCδ^flox/flox^ and PKCδ^flox/flox^ BMDM in both uninfected and infected states, rendering a segregated classification tree of LysM^cre^PKCδ^flox/flox^ and PKCδ^flox/flox^ samples on the horizontal axis of the heatmap (**Figure 4B-C**). In agreement with transcriptome, levels of Lyz2 was significantly decreased in the LysM^cre^PKCδ^flox/flox^ BMDM compared with PKCδ^flox/flox^ macrophages detected in both uninfected and infected total proteome clusters (**Figure 4B-C**). Interestingly, cumulative differentially expressed proteins (DEP) were associated with various catabolic processes in the uninfected macrophages suggesting baseline metabolic adaptation, while the regulation of neutrophil activation emerged as one of the prominent pathways during *Mtb* infection (**Figure 4D**). Additionally, phosphoproteomic dataset with stringent abundance and significance criteria identified a few phosphorylated increased and decreased proteins in BMDM lacking PKCδ at the uninfected timepoint (**Figure 4E**). However, only downregulated phospho-peptides achieved the significance and enrichment threshold during *Mtb* infection (**Figure 4F**). The association of phosphorylated DEP to Reactome pathways underscores critical phosphorylation events, including IRF-3-mediated type I IFN activation in the uninfected state, as evident in our transcriptome data (**Figure 4G**). RAC1, a small GTPase, is a key regulator of neutrophil recruitment to lung tissue, with its loss leading to reduced neutrophil influx and attenuated emphysematous lesions(*66*). During *Mtb* infection, we found activation of RAC1 was significantly increased as a consequence of PKCδ deletion in BMDMs, fortifying a crucial role of PKCδ in regulating neutrophil migration as reflected in our *in vivo* findings (**Figure 4G**). Given that several pathways involved in macrophage immune modulation during *Mtb* infection were influenced by the deletion of a single kinase, we next investigated whether the identified putative substrates or phosphorylated DEP could predict which other kinases likely to phosphorylate these substrates or DEP in uninfected and *Mtb* infected BMDM. By integrating phosphoproteomic data with the kinase-substrate prediction algorithm, Kinase Enrichment Analysis (KEA3)(*67*), we aimed to map upstream kinases to gain deeper insights into the dynamic kinase signaling events associated with PKCδ deficiency (**Figure 4H**). Surprisingly, we did not find PKCδ among other predicted kinases in uninfected state but PKCδ was apparent among the top 10 kinases in *Mtb* infection (**Figure 4H**), indicating the DEP that are dysregulated in macrophage immune-modulation, directly or indirectly interact with PKCδ, highlighting its role as a crucial regulatory kinase specifically during *Mtb* infection. Interestingly, a tyrosine kinase Fyn was amongst the highest-ranked kinases in both uninfected and infected phosphoproteome enrichment analyses. Overall, these results showed a wide spectrum of PKCδ involvement in macrophage signalling, including of neutrophil activation and migration during *Mtb* infection at the proteome level.

**Figure 4.**
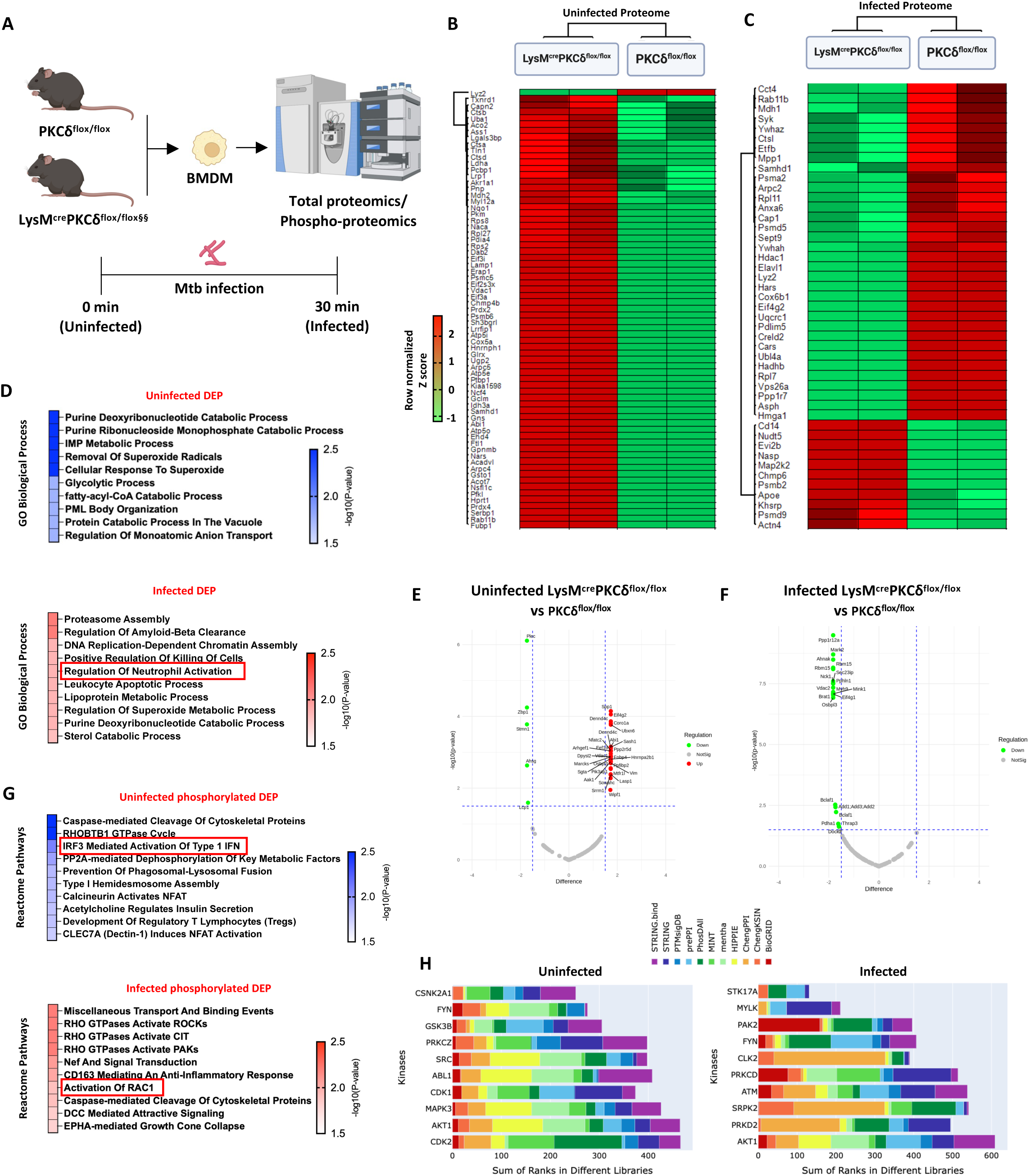
Total- & Phospho-proteome analysis revealed dysregulated protein clusters contributing to neutrophil activation in PKCδ deficient macrophages during *Mtb* infection. [A]. Experimental outline of Total- and Phospho-proteome analysis of bone-marrow-derived macrophages (BMDMs) from LysM^cre^PKCδ^flox/flox^ and PKCδ^flox/flox^ mice at indicated time points. **[B-C]** Heat-map analysis of total proteome of bone marrow-derived macrophages from LysM^cre^PKCδ^flox/flox^ and PKCδ^flox/flox^ mice at 0 minutes (Uninfected) and 30 minutes (MOI=1) post *Mtb* Infected (Infected) state. Protein clusters are arranged according to the similarity between their constituents. Horizontal and vertical clustering trees are depicting sample legends and differentially expressed proteins, respectively. **[D]** Gene Ontology (GO) analysis corresponding to differentially expressed proteins (DEP) shown in 0 min uninfected (B) and 30 min infected (C) group. The colour bars represents –log10(p-value) for the statistically significant top 10 biological processes. **[E-F]** Volcano plots for uninfected (0 min) and infected (30 min) phosphorylated DEP are shown between LysM^cre^PKCδ^flox/flox^ and PKCδ^flox/flox^ BMDMs. Upregulated, downregulated, and non-significant proteins are defined as red, green, and grey dots, respectively. Dotted line denotes threshold on X- and Y-axis. **[G]** Association of phosphorylated DEP to top 10 Reactome pathway database is shown at 0 min uninfected and 30 min infected group. **[H]** Kinase enrichment analysis of significant phosphorylated DEP corresponding to 0 min uninfected (E) and 30 min infected (F) group. Bars represent the mean rank of the top 10 kinases based on multiple library databases (color-coded above). Schematic cartoons by Biorender.com.

### PKCδ exerts intrinsic protection through independent GM-CSF and IFN signaling in macrophages against *Mtb* infection

In a TB-susceptible experimental mice model, elevated type I IFN signalling triggers pulmonary NETosis and promotes mycobacterial growth in the absence of GM-CSF(*68*). Signatures of dysregulated type I IFN signalling and enrichment of neutrophil activation and migration pathways led us to assess GM-CSF expression in LysM^cre^PKCδ^flox/flox^ BMDM during *Mtb* infection (**Supplementary Figure 5A**), consistent with the reduced GM-CSF levels *in vivo*. As colony-stimulating factor 2 receptor subunit alpha (CSF2RA), the receptor for GM-CSF, confers specificity to GM-CSF signalling(*69*), we also found a less prominent reduction in the expression of CSF2RA at 48 hpi in LysM^cre^PKCδ^flox/flox^ compared to PKCδ^flox/flox^ BMDM (**Supplementary Figure 5B**). This prompted us to assess whether exogenous GM-CSF counteracts the negative effects of elevated bacterial growth in LysM^cre^PKCδ^flox/flox^ BMDM. As expected, GM-CSF stimulated LysM^cre^PKCδ^flox/flox^ BMDM showed reduced mycobacterial growth compared to PKCδ^flox/flox^ BMDM (**Figure 5A**). Furthermore, GM-CSF stimulated LysM^cre^PKCδ^flox/flox^ BMDM also limited dysregulation of glycolysis and oxidative phosphorylation (OXPHOS) compared to PKCδ^flox/flox^ BMDM during *Mtb* infection, despite total ATP production still being higher in LysM^cre^PKCδ^flox/flox^ BMDM (**Figure 5B**). Additionally, the inflammatory status of LysM^cre^PKCδ^flox/flox^ BMDM was reversed as depicted by the M1 marker CD38, while M2 marker Egr2 remained reduced (**Figure 5C**). The inhibition of type I IFN signalling by Chlorpromazine (CPZ) lower the ability to produce type I IFN(*70*). We investigated whether blocking type I IFN signaling reduces the mycobacterial growth in LysM^cre^PKCδ^flox/flox^ BMDM and examined whether the addition of exogenous GM-CSF further enhances this reduction in mycobacterial burden. Surprisingly, we found a significant reduction of mycobacterial burden in CPZ-treated BMDM in LysM^cre^PKCδ^flox/flox^, which was restored but not significant with exogenous IFNβ (type I IFN) (**Figure 5D**). In contrast, CPZ and GM-CSF together were also able to suppress mycobacterial burden (**Figure 5D**), suggesting that exogenous GM-CSF dampens the burden independent of type I IFN signaling in the absence of PKCδ in macrophages, which requires further investigation.

**Figure 5.**
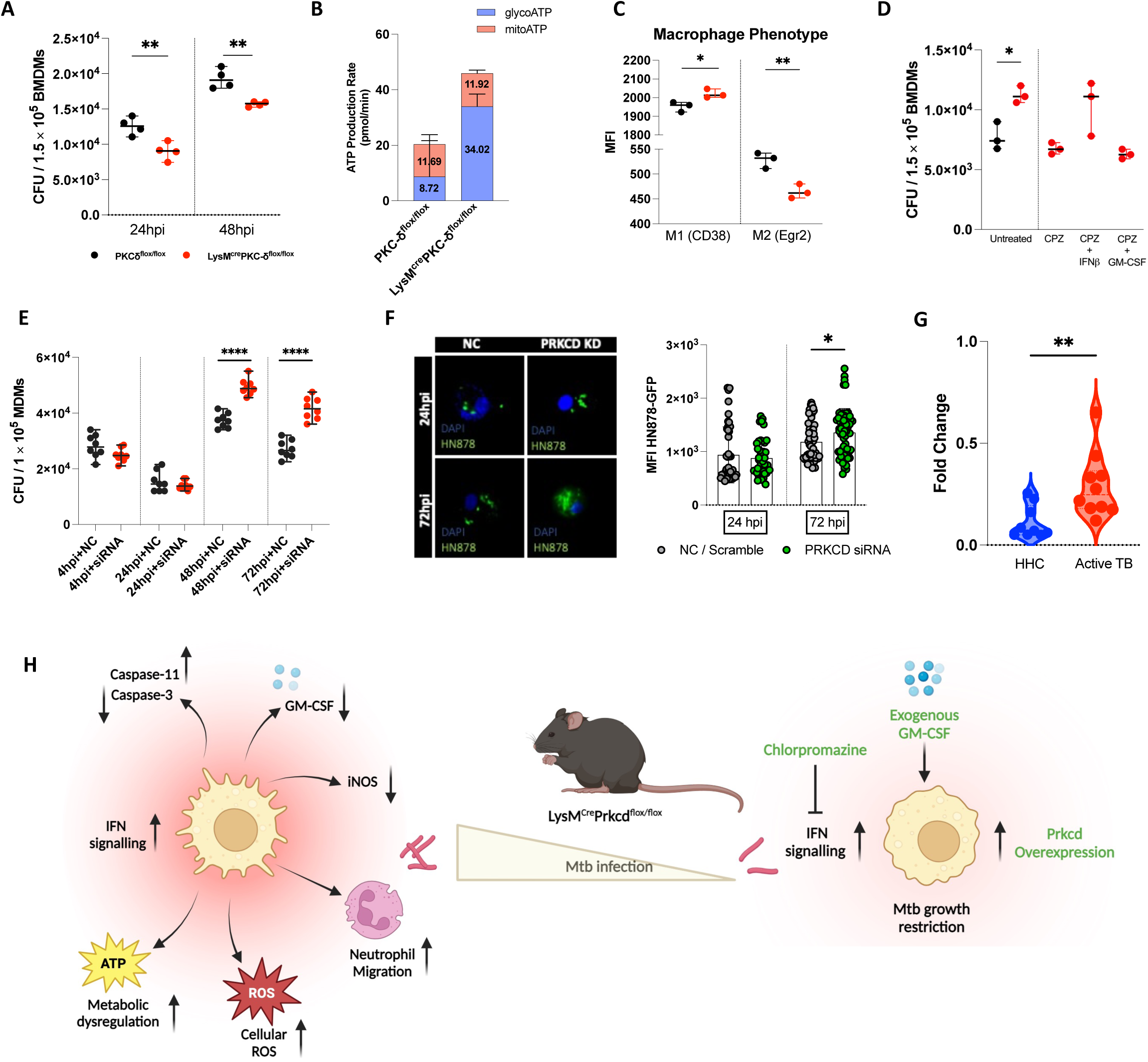
Exogenous GM-CSF and IFN signaling blockade independently restores the intrinsic protection in PKCδ deficient BMDMs against *Mtb* infection. [A]. Determination of mycobacterial burden in GM-CSF stimulated (50 ng/ml) bone-marrow-derived macrophages from LysM^cre^PKCδ^flox/flox^ and PKCδ^flox/flox^ mice during *Mtb* infection at indicated time points. **[B]** Total ATP production rate and metabolic dependence of GM-CSF stimulated bone-marrow-derived macrophages from LysM^cre^PKCδ^flox/flox^ and PKCδ^flox/flox^ mice at 24 hours post *Mtb* infection. **[C]** Determination of Mean Fluorescence Intensity (MFI) of macrophage phenotypic M1 and M2 marker by flow cytometry in GM-CSF stimulated bone-marrow-derived macrophages from LysM^cre^PKCδ^flox/flox^ and PKCδ^flox/flox^ mice at 24 hours post *Mtb* infection. **[D]** Determination of mycobacterial burden in CPZ (10 𝛍g/mL), IFN𝛃 (500U/mL), GM-CSF (50 ng/ml) stimulated bone-marrow-derived macrophages from LysM^cre^PKCδ^flox/flox^ at 24 hours post Mtb infection. **[E]** Determination of mycobacterial burden in the scramble siRNA treated and PRKCD deficient human monocyte-derived macrophages at indicated time points. **[F]** Comparison between the mean fluorescence intensity (MFI) of HN878-GFP at 24 hpi and 72 hpi using Fiji (Image processing module). **[G]** mRNA expression of PRKCD in bronchoalveolar lavage (BAL) samples acquired from active TB patients and household contacts (HHC) (N= 10/arm). **[H]** Proposed model of PKCδ in regulation of immunomodulatory macrophage functions during Mtb infection. All data shown mean+/- SD and is representative of two independent experiments performed in triplicates or quadruplicates. Statistical analyses were performed using an unpaired student t-test. Asterisks are defining significance compared to the control group as: *p < 0.05, **p < 0.01, ****p < 0.0001. Schematic cartoons by Biorender.com.

To further assess the relevance of our findings in human macrophages, first, we assessed the expression kinetics of human PRKCD gene in monocyte-derived macrophages (MDM) during *Mtb* infection. We found that the mRNA expression of PRKCD increased significantly from 4 hpi to 24 hpi remaining stable thereafter (**Supplementary Figure 5C**), consistent with Prkcd expression in mouse macrophages(*16, 52*). Of interest, we successfully silenced PRKCD using anti-PRKCD siRNA and achieved a knockdown efficiency of ∼90% and above in MDM (**Supplementary Figure 5D**). We then infected PRKCD-silenced MDM and observed an increase in mycobacterial growth at 48 and 72 hpi in comparison to MDM that was transfected with scramble siRNA (**Figure 5E**). This was corroborated by imaging where we detected an increase in the bacterial growth, as measured by mean-fluorescence intensity (MFI) of GFP-expressing *Mtb* HN878 bacilli in PRKCD-silenced MDM compared to scramble siRNA-treated MDM at 72 hpi (**Figure 5F**). As *Mtb* infection induces robust immunological responses in the absence of PKCδ in macrophages, we explored whether its expression changes during the transition from infection to active TB disease. We analysed PRKCD gene expression in bronchoalveolar lavage (BAL) samples obtained from active TB patients, as it exhibits the most favourable clinical manifestation of TB disease(*71*). Expectedly, PRKCD mRNA expression was significantly elevated in the active TB group compared to the household contact control group (**Figure 5G**), establishing a strong basis for further exploration of PRKCD as a central regulator of immunoregulatory functions in clinical settings during *Mtb* infection. Systemic evaluation of PKCδ in macrophage immune perturbations showed an undiscovered role in balancing *Mtb* infection outcome on one hand, by modulating antimicrobial effector functions compelling the bacterial growth, and on the other hand, augmented bacterial growth was reduced by addition or blocking of PKCδ mediated immune correlates (**Figure 5H**).

## Discussion

Based on findings in PKCδ null mice (PKCδ^-/-^) and PKCδ deficient individuals in biological and functional studies in different disease settings, it is evident that PKCδ is crucial to maintain immune homeostasis against microbial intruders and autoimmune disorders(*14–16, 72, 73*). Although, PKCδ^-/-^ mice were characterized by differences in lymphocytes at naive state(*18, 74*), very limited studies have been conducted for myeloid-specific deficiency of PKCδ and its immunomodulatory characteristics *in vivo*. As the tissue environment is governed by the development, activation, recruitment, and function of macrophages both in naive and disease states, we therefore generated macrophage-specific knockout mice (LysM^cre^PKCδ^flox/flox^) and characterized in light of the previous abnormalities observed in PKCδ^-/-^ models. Immune system-focused characterisation under homeostatic conditions revealed no significant imbalance in the LysM^cre^PKCδ^flox/flox^ model.

In this study, we investigated the immunomodulatory functions of macrophage-specific deletion of PKCδ during *Mtb* infection *in vivo*. Our findings revealed an increased mycobacterial burden accompanied by exacerbated lung pathology in the LysM^cre^PKCδ^flox/flox^ mice, thereby confirming a critical role of macrophage-expressed PKCδ in host defense. These results suggest that the heightened susceptibility observed in our previously reported PKCδ^-/-^ mice was predominantly driven by PKCδ in macrophages. Host defense mechanisms depend on a finely tuned balance; disruption of the equilibrium could favor *Mtb* proliferation, resulting in disease progression, and/or trigger tissue destruction through excessive immune activation(*75*). Evidently, the absence of PKCδ in macrophages rendered decreased reactive nitrogen intermediates defined as reduced iNOS at the early stages of *Mtb* infection in the lungs, promoting pathogen-permissive environment detrimental to the host(*76*). The progression of TB pathogenesis follows a temporal relationship wherein logarithmic growth of the pathogen coincides with the gradual development of *Mtb* antigen-specific T cell responses, which engage infected macrophages to mediate host defense(*24*). CD4^+^/CD8^+^ T cells are pivotal to control *Mtb* in both humans and mice (*77*). The conventional T cell frequencies were not significantly affected by the ablation of PKCδ in macrophages, surprisingly, T cell memory and effector subsets were increased at the early stage of *Mtb* infection. T cells are required for the control of infection *in vivo*; however, these immune cells cannot eliminate the pathogen or protect the host from a fatal infection alone(*77, 78*), which may explain the growth of the pathogen in the presence of robust T memory immune response in the LysM^cre^PKCδ^flox/flox^ mice in the acute stage. However, this response has overcome by *Mtb* over time, leading to dissemination to the spleen in the chronic stage of *Mtb* infection. Prolonged challenge with *Mtb* antigen has been shown to facilitate reduction in IL-2 and CD8^+^ T memory cells *in vivo*, resulting in impaired antibacterial immunity(*24, 77*). We found a significant reduction in the IL-2 levels with restored effector and memory T cell turnover in the LysM^cre^PKCδ^flox/flox^ mice similar to the control animals at the chronic stage of *Mtb* infection. This varied T cell interplay modulating IL-2 levels at various stages of *Mtb* infection in the absence of PKCδ in macrophages is critical and needs further evaluation. *Mtb* can also disrupt innate immunity processes by subverting various myeloid cells. Recent studies have explained a nutrient-replete niche that supports bacterial growth and tissue damage by neutrophilic inflammation during *Mtb* infection(*76*). Moreover, NOS2^-/-^ mice exhibited an enhanced neutrophilic influx into the lungs, promoting increased susceptibility to TB(*76*). This accords with our observation that neutrophils in the lungs of the LysM^cre^PKCδ^flox/flox^ mice produce fewer nitrogen intermediates in response to acute *Mtb* infection. Among the major myeloid cells at the site of infection, macrophages play a key role both in disease control as well as progression(*39*). Studies have established the restrictive nature of interstitial macrophages (IM) exhibiting a distinct metabolic state by contrast with permissive alveolar macrophages(*39*). We inferred that increased mycobacterial burden was likely associated with reduced frequencies of IM in the lungs of LysM^cre^PKCδ^flox/flox^ mice. Also, murine IM subsets are known to produce IL-10 among macrophage/monocyte lineages(*79*). Therefore, reduced IL-10 levels observed in the lungs of the LysM^cre^PKCδ^flox/flox^ mice support this immunophenotypic outcome of depleting inflammatory macrophages during *Mtb* infection.

The phagocytosis of *Mtb* by macrophages in the lungs initiates host protection from infection and involves various cellular mechanisms(*80, 81*). We observed switching of various cell death modalities in the absence of PKCδ in macrophages during *Mtb* infection. Cellular ROS has been described as a bactericidal effector function during pathogen engulfment by macrophages(*82*). Surprisingly, we found a significant increase in the cellular ROS levels in the LysM^cre^PKCδ^flox/flox^ macrophages, despite the enhanced ROS not correlating with improved control of *Mtb* infection. This is unexpected, as ROS are generally believed to exert antimicrobial effect against infective agents. However, our findings raise important question of whether ROS on this context function primarily as antimicrobial agents or inadvertently favor *Mtb* survival. Mice lacking NOX2, a prominent ROS-generating transmembrane protein, are less susceptible to *Mtb* infection(*83*) and excessive or incorrectly localized ROS can be detrimental, leading to host damage(*82*). LysM^cre^PKCδ^flox/flox^ macrophages had higher mitochondrial ROS levels, indicating that PKCδ may regulate excessive ROS-mediated inflammation following *Mtb* infection *in vivo*. We also detected less expression of iNOS and NO, suggesting that PKCδ is essential for NO-mediated control of enhanced bacterial growth in macrophages. We probed additional antimicrobial effector mechanisms and PKCδ emerged as a key player in caspase-3-mediated apoptosis that drives infected macrophage killing as well as the MHCII-mediated antigen presentation during *Mtb* infection *in vitro*. Furthermore, previous study suggests a link between elevated mitochondrial ROS and enhanced IL-1β production via activation of the NLRP3 inflammasome(*84*). The role of IL-1β is dichotomous in *Mtb* infection, with some studies suggesting a role in host protection whilst others show the host vulnerability in sustained IL-1β exposure(*85*). Additionally, a recent study proposed that pharmacological and genetic suppression of NLRP3 inflammasome would decrease bacterial viability in macrophages for *Mtb* strains that produce a strong IL-1β response(*86*). Based on our observations of ROS elevation, we have found an increase in IL-1β secretion in LysM^cre^PKCδ^flox/flox^ macrophages, which in turn was associated with the modest upregulation of NLRP3 and AIM2 inflammasome genes. We observed a modest yet previously unrecognized immune response signature suggestive of caspase-11 mediated non-canonical inflammasome activation in PKCδ-deficient macrophages in *Mtb* infection. Although the precise mechanism is unclear, the detrimental role of excessive production of IL-1β may augment bacterial growth in the context of PKCδ deficiency in macrophages following *Mtb* infection, which needs further exploration. Numerous investigations have demonstrated that NLRP3-mediated increased IL-1β production during inflammation is ATP-dependent(*87, 88*). This was evident in our results with increased ATP production in LysM^cre^PKCδ^flox/flox^ macrophages, functionally implicating a role in bacterial persistence in the absence of PKCδ in murine macrophages during *Mtb* infection. Metabolic profiling further revealed this enhanced ATP generation was primarily driven by glycolysis, however, redundant towards OXPHOS. Notably, Mills and colleagues have described an LPS-induced rise in glycolytic activity that generates sufficient cytosolic ATP reducing the need for mitochondrial oxidative phosphorylation. This leads to build-up of mitochondrial membrane potential and allowing Complex I to produce pro-inflammatory mitochondrial ROS through reverse electron transport(*89*).

We found that absence of PKCδ also perturbed the expression of several genes associated with macrophage antimicrobial effector functions during *Mtb* infection. Transcriptomic analysis further indicated that the differentially expressed genes were linked to type I IFN signalling in the absence of PKCδ in the naive state, a pattern that persisted during *Mtb* infection as well, albeit, to a lesser extent. Constitutively, this likely contributes to a less inflammatory macrophage profile and the accumulation of permissive myeloid cells, promoting impaired bacterial control as reported in previous studies(*90, 91*). Also, exacerbation of *Mtb* infection due to type I IFN signalling, which leads to neutrophilic recruitment, has been reported(*59, 68*). The absence of PKCδ displayed a similar pattern of upregulated neutrophil associated genes following hyperactivation of type I IFN signalling, positioning PKCδ as an upstream kinase regulator of the type I IFN signalling pathway, which contributes to intensified *Mtb* infection of macrophages. Comprehensive proteome analysis also demonstrated that the absence of PKCδ is possibly modulating RAC1-driven neutrophil activation in *Mtb* infection. A critical role of RAC1 as a physiological integrator for neutrophil chemotaxis(*66*) and migration into the lung during inflammation has been well documented(*92*). However, the association of RAC1 activation in murine macrophages support an unexplored indirect role of PKCδ in regulating neutrophil migration during *Mtb* infection. Additionally, GM-CSF has also been well-documented in terms of its role in mediating *Mtb* control and inflammation *in vivo*(*93, 94*). Despite no observable changes in the frequencies of alveolar macrophages, GM-CSF levels were significantly reduced in the lungs during the disease progression. This is particularly noteworthy, as previous studies established a clear link between GM-CSF and bactericidal activity of alveolar macrophages(*95*). While GM-CSF can be produced by a variety of host immune cells such as macrophages, conventional/ non-conventional T cells, and alveolar epithelial cells(*32*), we were limited in able to determine the major sources of GM-CSF in the absence of PKCδ in macrophages *in vivo*. However, the macrophages could be amongst the cells which are a major producer of GM-CSF since the latter was abolished in the absence of PKCδ *in vitro*. Consistent with our findings, Mishra *et al* reported that macrophages from active TB patients exhibited lower levels of GM-CSF(*96*), aligning with our *in vitro* observation of diminished production of GM-CSF in the absence of PKCδ in murine macrophages during *Mtb* infection. While others have reported an impaired GM-CSF signalling leads to type I IFN induced neutrophil migration into lungs during murine *Mtb* infection(*68*), we found an unexplored link, showing that this effect might be facilitated by PKCδ in murine macrophages. Interestingly, exogenous GM-CSF supplementation in LysM^cre^PKCδ^flox/flox^ macrophages reduced mycobacterial growth with a subsequent restriction of uncontrolled metabolic reprogramming during infection and restored the inflammatory macrophage status. Similarly, pharmacological blockade of type I IFN signalling by Chlorpromazine (CPZ) reduced PKCδ-deficiency mediated augmented *Mtb* burden, which was also sustained with the addition of exogenous GM-CSF. GM-CSF licenses signaling through IFNyR (type II IFN) to confer monocytic inflammatory functions(*97*). PKCδ-deficient macrophages also confirmed GM-CSF licensing may not be dependent on type I IFN transcriptional signalling to exhibit inflammatory response, which was blocked by CPZ treatment.

Previously, whole blood transcript profiling showed a significant increased mRNA expression of PKCδ in active TB patients compared to healthy and LTBI individuals as well as PRKCD protein in human caseous and necrotic lung granulomas(*16*). We also observed higher mRNA expression of PKCδ in BAL samples from active TB patients compared to household contacts, pointing towards the involvement of PKCδ kinase at the site of infection and presenting a prospective target for future therapeutic intervention. Upon employing human MDM *Mtb* infection approach, we observed a comparable increase in PKCδ signature, indicating a consistent activation pattern in response to infection. Furthermore, conventional knockdown of PKCδ in MDMs showed an increase in mycobacterial growth, highlighting PKCδ is also optimal for macrophage restriction of *Mtb* infection in humans. Our study suggests that as a key hub of intracellular immune modulators – PKCδ acts on various cellular modalities which restrict bacterial growth in macrophages and inflammation *in vivo*. The protective effect was also enhanced by overexpressing PKCδ, exogenous addition or blocking of PKCδ regulated upstream and downstream immune correlates. Due to the broad role of PKCδ in immune modulation, our study confers that GM-CSF represents one of the mechanism(s) through which PKCδ attenuated immune dysregulation during *Mtb* infection. Moreover, sustained suppression of PKCδ could potentially trigger direct or indirect compensatory mechanisms via other family members of PKC family potentially driven by macrophage plasticity in response to *Mtb* infection. This underscores the need for focused investigation to validate PKCδ as a viable therapeutic target, mainly through the development of highly specific pharmacological activators to harness its immunoregulatory potential without invoking broader, possibly maladaptive pathways. In conclusion, our study reveals that PKCδ is a key hub governing macrophage immunodynamics during *Mtb* infection.

## Materials and Methods

### Mouse Strains

To generate the macrophage-specific PKCδ knockout mice, we crossed loxP flanked PKCδ mice of C57BL/6 strain with in-house LysM^cre^ mice of C57BL/6 strain for three generations to achieve homozygous knockout model LysM^cre^PKCδ^flox/flox^. LoxP flanked PKCδ (PKCδ^flox/flox^) mice were gifted from Prof. C Ronald Kahn at the Joslin Diabetes Center, USA and considered as a littermate control for all downstream experiments(*27*). All mice were generated and maintained under specified pathogen-free conditions by the Animal Research Facility under strict guidelines of approved protocols by the Animal Research Ethics Committee (AREC), University of Cape Town. Experimental mice were matched for age (8-12 weeks) and sex. All procedures were conducted in the Research Animal Facility (RAF) Biosafety Level 3 (BSL3).

### Mouse genotyping

Genomic DNA was extracted from both PKCδ^flox/flox^ and LysMcrePKCδ^flox/flox^ mice tails to confirm the genotype. Multiplex PCR on T100™ Thermal Cycler (Bio-Rad) was performed to amplify the transcripts using the following primers: PKCδ: Forward sequence: 5’- CTG GGT AAC TTA ACA AGA CC-3’; Reverse sequence: 5’- CTG CTA AAT AAC ATG TTC GGT CC-3’; LysM^cre^: Forward sequence: 5’- CCC AGA AAT GCC AGA TTA CG-3’, Reverse sequence: 5’-CTT GGG CTG CCA GAA TTT CTC-3’. Optimal PCR conditions include an initial denaturation (94°C for 3 min), denaturation (35 cycles; 94°C for 30 sec), annealing (60°C for 30 sec), extension (72°C for 30 sec) and final extension (72°C for 5 min) were conducted. The PCR product was separated on 1.6% Agarose gel (SYBR Safe DNA gel stain, Thermofisher) and visualized under UV light on Syngene G: Box and analyzed on GeneSnap (Syngene, Cambridge). Expected PCR amplicon sizes: Wildtype(n/n): 350bp; floxed PKCδ (fl/fl): 425bp; LysM^cre^PKCδ^flox/flox^ (LysM^cre+^): 700bp.

### Ethical considerations

All experimental animals were operated following the Animal Research Ethics Committee (AREC) approved protocols (AREC Permit No: 019/023 and 019/031 and 022/024) and regulations under the South African National Standard (SANS 10386:2008) at the Faculty of Health Sciences, University of Cape Town. Synthetic complementary DNA were obtained from deidentified biobanked BAL samples from completed studies previously approved by the Health Research Ethics Committee of Stellenbosch University (N13/05/064 and N10/08/276).

### Generation of bone-marrow-derived macrophages

As previously described(*52*), bone-marrow-derived macrophages (BMDM) were generated from 8-12 weeks old PKCδ^flox/flox^ and LysM^cre^PKCδ^flox/flox^ mice. Briefly, bone marrow cells were flushed out from femurs and cultured for 10 days at 37°C under 5% CO2 in sterile tissue culture grade Petri dishes (140mm x 20mm Petri dish, Nunc, Denmark) consisting of PLUTZNIK media (DMEM containing 30% L929 cell-conditioned medium, 10% fetal calf serum, 5% horse serum, 1 mM sodium pyruvate, 2 mM L-glutamine, 0.1 mM β-mercaptoethanol, 100 U/ml penicillin G, and 100 μg/ml streptomycin. BMDM were harvested and plated in desired culture plates in complete media (DMEM with 10% fetal calf serum) to proceed with the downstream experimental procedure.

### Quantitative real-time polymerase chain reaction (qRT-PCR)

RNA was extracted using the RNeasy Mini Kit (Qiagen, cat# 74106) from stored RNA samples in 350 μl of RLT lysis buffer with 3.5 μl β-mercaptoethanol. 400ng normalized RNA was reverse transcribed using a High-Capacity cDNA Reverse Transcription Kit (Applied Biosystems, cat# 4368814) with random primers to yield first-strand cDNA following the manufacturer’s protocol. Desired gene expressions were amplified using Fast SYBR™ Green Master Mix (Applied Biosystems, cat# 4385612) and analyzed using Quantstudio 7 (Applied Biosystems, USA). The qRT-PCR conditions were as follows: Stage 1 (x1 cycle): Pre-incubation 95°C for 10 min; Stage 2 (x45 cycle): Denaturation 95°C for 15 sec, Annealing 60°C for 10 sec, Extension 72°C for 15 sec, Final acquisition 80°C for 1 sec. Candidate gene expressions were normalized using an endogenous housekeeping control Hprt. Primer sequences are listed in ***Table S1*** with their respective gene accession number.

### Western blot analysis

Sodium dodecyl sulfate-polyacrylamide gel electrophoresis (SDS-PAGE) and Western Blot analysis were performed as previously described(*98*). Briefly, bone-marrow-derived macrophages from PKCδ^flox/flox^, LysM^cre^PKCδ^flox/flox^, PKCδ overexpressed and control RAW264.7 murine macrophage cells (3 x 10^6^) were seeded in a 6-well tissue culture graded plate containing complete media (DMEM containing 10% fetal calf serum) for overnight at 37°C under 5% CO2 incubator. Cells were washed and lysed with ice-cold RIPA buffer containing protease and phosphatase inhibitors for 30 min at 4°C. BCA Protein Assay Kit (Thermofisher Scientific Pierce™, cat# 23225) was used to determine the total protein concentration. An equal amount (20 μg) of protein were boiled at 100°C for 5 min with 1X loading dye (2% SDS, 5% 2-mercaptoethanol, 10% glycerol, 0.002% bromophenol blue, 0.62 M Tris-HCl, pH 6.8). Samples were then electrophoresed on 12% SDS-PAGE gel (Mini-PROTEAN® system, Bio-Rad) and transferred to nitrocellulose membrane (Sigma) using the Mini Trans-Blot® Cell system (Bio-Rad). Membrane blocking was performed for 2 hours on a shaker at room temperature with 5% w/v BSA, 1X TBS (20 mM Tris with 150 mM NaCl), and 0.1% Tween20 (blocking buffer). Next, the membrane was probed with recombinant anti-PKC delta antibody (Abcam; ab182126) or GAPDH [Santa Cruz Biotechnology; (sc47724)] primary antibodies (in 5% BSA blocking buffer; 1:1000 dilution) according to the manufacturer’s protocol at 4°C overnight and goat anti-rabbit IgG H&L (Abcam; ab97040) secondary antibody (in 5% BSA blocking buffer; 1:10000 dilution) at room temperature for one hour. Immunoblots were developed using the KPL LumiGLO® Reserve Chemiluminescent Substrate Kit (SeraCare Life Sciences; cat# 5430-0042(54-61-02)) on the iBright FL1000 Imaging System (Thermofisher Scientific).

### *Mtb* infection and determination of mycobacterial burden in vivo

A virulent strain of *Mtb* (HN878) was grown in complete 7H9 media to log phase and glycerol stocks were made to infect mice by an intranasal administration procedure as previously described(*52*). Stock solutions of *Mtb* were thawed and washed once with phosphate-buffered saline (PBS) to eliminate glycerol before infection. The inoculum was prepared in sterile saline containing 0.05% Tween-80. Anesthetized mice were challenged with 50μl of *Mtb* inoculum (25μl/nostril). Inoculum dose uptake was also confirmed one day post-infection by determining the mycobacterial burden in the lungs of 4-5 infected mice. Mycobacterial load in the lungs and spleen of *Mtb* infected mice was determined at the indicated time points. Briefly, organs were aseptically collected and weighed from euthanized mice and homogenized in 0.04% Tween-80. Organ homogenates were subjected to plated as 10-fold dilution for CFU growth on Middlebrook 7H10 (BD Biosciences) agar plates supplemented with 10% OADC and 0.5% glycerol. Agar plates were incubated for 14-21 days at 37°C before colonies were counted.

### Lung histopathology and immunohistochemistry

The right superior lobe of the lung from *Mtb* infected mice was collected in formalin solution (10% formaldehyde in 1X PBS) and processed for cryo-sectioning with the Leica TP 1020 benchtop processor following embedded in paraffin wax. Processed sections were cut using Leica Sliding Microtome 2000R in four 3 μm thick sections for H&E staining and two 3 μm sections for iNOS staining. Stained sections were scanned at 40X in the Olympus VS120 virtual microscope for image acquisition. Alveolar space and iNOS^+^ areas were measured using Qupath v3.0.2 open-source software. Total alveolar spaces were quantified by subtracting the H&E positive area from the total lung tissue area. Similarly, the iNOS^+^ area was calculated as the area positively stained with diaminobenzidine (DAB) stain.

### Immune cell population in lungs and spleen by fluorescence-activated cell sorting

Left lobes of the lung were collected and digested in lung digestion buffer (DNase1, Collagenase type 1, DMEM) at 37°C for one hour to prepare a single-cell suspension. Digested lungs were passed through 100 μm and 70 μm sterile cell strainers (SPL Life Sciences) whereas spleens were mechanically digested through 70 μm and 40 μm sterile cell strainers (SPL Life Sciences). Cells were washed with complete media (DMEM + 10%FCS) following red cell lysis buffer (150 mM NaCl, 10 mM KHCO3 and 0.1mM Na2EDTA) incubation for 5 min at room temperature. Then, cells were washed again with complete media before being counted with CytoSMART (Corning) automated cell counter and seeded at approximately 1 x 10^6^ cells per well in a 96-well U-bottom (Corning) plate. Seeded cells were washed once with PBS and stained with dead cell marker (575V Viability Dye, BD Biosciences) for 15 min at room temperature. After that cells were washed with FACS buffer (0.5% BSA, 0.5% Sodium azide (NaN3), 1x PBS) and surface stained for the following lymphoid and myeloid-specific markers: Lymphoid markers (Lung) - Gamma delta (γδ) T cells: CD3+ γδTCR+; Natural Killer (NK) cells: NK1.1^+^CD3^-^; B cells: CD19^+^CD3^-^; T cells: CD3^+^CD19^-^; CD4 T cells: CD3^+^CD4^+^; CD8 T cells: CD3^+^CD8^+^, CD4^+^ Naive: CD3^+^CD4^+^CD62L^+^CD44^-^; CD4^+^ Central Memory: CD3^+^CD4^+^CD62L^+^CD44^+^; CD4^+^ Effector/Effector Memory: CD3^+^CD4^+^CD44^+^CD62L^-^; CD4^+^ Double Negative: CD3^+^CD4^+^CD44^-^CD62L^-^; CD8^+^ Naive = CD3^+^CD8^+^CD62L^+^CD44^-^; CD8^+^ Central Memory = CD3^+^CD8^+^CD62L^+^CD44^+^ ; CD8^+^ Effector/Effector Memory = CD3^+^CD8^+^CD44^+^CD62L^-^; CD8^+^ Double Negative: CD3^+^CD4^+^CD44^-^CD62L^-^; CD4^+^ Exhaustion Memory: CD3^+^CD4^+^PD1^-^KLRG1^+^; CD8^+^ Exhaustion Memory (lung): CD3^+^CD8^+^PD1^-^ KLRG1^+^; CD8^+^ Tissue Resident Memory: CD3^+^CD8^+^CD103^+^CD69^+^; CD8^+^ Memory Precursor: CD3^+^CD8^+^KLRG1^-^CD127^+^; CD8^+^ Short-lived Effector: CD3^+^CD8^+^KLRG1^+^CD127^-^; Myeloid markers (Lung) - Neutrophils: Ly6G^+^CD11b^+^; Eosinophils: SiglecF^+^CD11b^+^CD64^-^; Dendritic cells (DC): CD11b^+^CD11c^+^MHCII^+^; Monocytes: Ly6G^+^CD11b^+^CD64^-^; Alveolar macrophages: CD64^+^MerTK^+^SiglecF^+^CD11c^+^; Interstitial macrophages: CD64^+^MerTK^+^SiglecF^+^CD11b^+^CD11c^-^; Myeloid markers (Spleen) - Neutrophils: Ly6G^+^CD11b^+^; Monocytes: CD11b^+^Ly6C^+^CD11c^-^; Marginal zone macrophages: CD11b^high^F4/80^low^CD11c^-^; Red pulp macrophages: F4/80^high^CD11b^low^CD11c^-^; CD11b^+^ DC = CD11c^+^MHCII^+^CD11b^+^. Surface-stained cells were acquired using BD LSR Fortessa (BD Biosciences Immunohistochemistry Systems) and data was analysed by FlowJo v10.6.1 software (TreeStar, US). All stained cells positive for specific markers are calculated as a percentage of live cells.

### Cytokine, chemokine, and growth factor profiling in lung homogenates

Lung homogenates were spun at 3000 g for 5 min to procure cell-free supernatant analysed using standard sandwich ELISA assay. Coating, standard, and detection antibodies were obtained from BD Biosciences, Biolegend, and R&D Scientific. The assay was performed to detect different cytokines (TH1: IL-1α, IL-1β, IL-2, IL-6, IL-12p40, IL-12p60, IFN-β, IFN-γ; TH2: IL-4, IL-5, IL-10, IL-13, TGF-β; TH17: IL-17, IL-23), chemokines (CCL3, CCL5, CXCL1, CXCL2, CXCL5, CXCL10) and growth factors (IGF-1, GM-CSF) levels according to manufacturer’s dilutions and protocol. TMB microwell peroxidase substrate (KPL International) for streptavidin-HRP conjugates or 1 mg/ml p-nitrophenyl phosphate disodium salt hexahydrate (Sigma, cat# N2765) for streptavidin-AP conjugates. VersaMax™ microplate spectrophotometer (Molecular Devices, Sunnyvale, California) was used to measure optical density.

### Lentiviral overexpression

The RAW264.7 murine macrophage cell line was obtained from ATCC, cultured, passaged, and seeded at 0.5x10^5^ cells in a tissue culture grade 24-well plate for following lentivirus-packaged ORF clone of PKCδ (MR225184L4V, Origene) transduction according to the manufacturer’s protocol. Briefly, frozen lentivirus particles were thawed and added to the cells with a fresh culture medium (DMEM with 10% fetal calf serum) of 500μl at a multiplicity of infection (MOI=5) with 8 μg/ml of Polybrene (Sigma). Cells were incubated with the lentivirus-contained medium for 18-20 hours at 37°C and replaced with a fresh medium, followed by the stable cell selection with 2.5 μg/ml Puromycin (ThermoFisher) for 10-14 days. PKCδ overexpressed cells were detected using a ZOE fluorescence microscope (BioRad) for GFP expression, followed by the collection of RNA and protein to confirm overexpression confirmation by qRT-PCR and western blot, respectively, and downstream *Mtb* infection experiment.

### *Mtb* infection and determination of mycobacterial burden in vitro

A virulent strain of *Mtb* (HN878) was grown in complete 7H9 media to log phase and glycerol stocks were made for *in vitro* infection of the BMDM and MDM as previously described(*99*). Stock solutions of *Mtb* were thawed and washed once with 1X PBS to get rid of glycerol before infection. Seeded BMDM and MDM were infected at multiplicity of infection (MOI) 1 and 5, respectively. At indicated time points, cells were washed with 1X PBS, lysed with 0.1% Triton X-100, followed by 10- and 100-fold dilution, and plated on Middlebrook 7H10 (BD Biosciences) agar plates supplemented with 10% OADC and 0.5% glycerol. Agar plates were incubated for 14-21 days at 37°C before colonies were counted. RNA and protein were collected at indicated time points from *Mtb* infected cells for downstream processing. A similar procedure was followed to enumerate mycobacterial load in PKCδ overexpressed and control RAW264.7 murine macrophages.

### Enzyme-Linked Immunosorbent Assay (ELISA) and Griess assay

BMDM were seeded in a 96-well cell culture plate and infected with *Mtb* (MOI=1) at indicated time points, followed by the procurement of cell-free supernatant analysed using a standard sandwich ELISA assay. Coating, standard, and detection antibodies were obtained from BD Biosciences, Biolegend, and R&D Scientific. The assay was performed to detect proinflammatory cytokines: IL-1𝛂, IL-1𝛃 and IL-6 according to the manufacturer’s dilutions and protocol. TMB microwell peroxidase substrate (KPL International) for streptavidin-HRP conjugates or 1 mg/ml p-nitrophenyl phosphate disodium salt hexahydrate (Sigma, cat# N2765) for streptavidin-AP conjugates. Nitrite levels in *Mtb* infected cell-free supernatants were measured using the Griess reagent assay. Briefly, cell supernatants were incubated with 1% sulfanilamide in 2.5% phosphoric acid for 10 minutes at room temperature in the dark, followed by 0.1% naphthyl-ethylene-diamine in 2.5% phosphoric acid for another 10 minutes. VersaMax™ microplate spectrophotometer (Molecular Devices, Sunnyvale, California) was used to measure optical densities at 405 nm for AP-conjugates and 450 nm for HRP-conjugates.

### Cellular and mitochondrial Reactive Oxygen Species (ROS) assay

BMDM were seeded in a 96-well cell culture plate and infected with *Mtb* (MOI=1) at indicated time points, followed by incubation with either 5 μM CellROX Green Reagent (ThermoFisher) or 100 nM MitoTracker Red CM-H2XRos (ThermoFisher) for 30 minutes at 37°C. Cells were then washed with 1X PBS three times. Wells with only media were used as blank. Fluorescence was measured on Spectramax iD3 multi-mode reader (Molecular Devices) with the excitation of 485 nm and emission of 525 nm for CellROX and excitation of 579 nm and emission of 599 nm for MitoTracker intensity.

### pHrodo labelling of *Mtb* and detection of phagosome maturation

*Mtb* stocks were thawed and labelled with 100 nM pHrodo iFL Green STP ester, amine-reactive dye (TheroFisher) in 100 mM sodium bicarbonate buffer (pH 8.5) at room temperature for 30 minutes. Labelled *Mtb* the washed with 1X PBS three times before infection of BMDM. Phagolysosomal pH-mediated fluorescence was recorded at indicated time points with excitation of 500 nm and emission of 540 nm on Spectramax iD3 multi-mode reader (Molecular Devices).

### Determination of macrophage apoptosis, activation, and polarization by flow cytometry

*Mtb* infected BMDM were washed once with 1X PBS and stained with dead cell marker (575V Viability Dye, BD Biosciences) for 15 min at room temperature. After that cells were washed with FACS buffer (0.5% BSA, 0.5% Sodium azide (NaN3), 1x PBS) and surface stained for the following surface markers: CD11b – PerCPCy5.5 (Clone M1/70), F4/80 – PeCy7 (Clone BM8), CD80 – BV421 (Clone 16-10A1), CD86 – AF700 (GL-1), MHCII – APC (Clone M5/114.15.2), CD38 – FITC (Clone H-11) and Egr2 – APC (Clone erongr2). Surface-stained cells were permeabilized and blocked (Permeabilization buffer + 2% IRS + 1% 2.4G2 FcyRII/III blocker) at 4°C for 10 minutes, followed by the staining for intracellular marker: active Caspase-3 – PE (Clone 51-68655X) at 4°C for 30 minutes. Intracellularly stained cells were resuspended in FACS buffer and acquired using BD LSR Fortessa (BD Biosciences Immunohistochemistry Systems) and data was analysed by FlowJo v10.6.1 software (TreeStar, US). All stained cells positive for specific markers are calculated as a percentage of live cells.

### Seahorse XF real-time ATP rate assay

BMDM (5 x 10^4^) were seeded in Seahorse XFp cell culture mini plates before *Mtb* infection. Cell culture medium was removed, washed with pre-warmed Seahorse assay media (10 mL of Seahorse XF DMEM Medium, pH 7.4 with 10 mM of XF glucose, 1 mM of XF pyruvate, 2 mM of XF glutamine), and replaced with *Mtb* (MOI=1) containing assay media to the total volume of 50 μl per well. Seahorse XF real-time ATP rate assay sensor cartridge and utility plate were subjected to overnight hydration and a 1-hour calibration period in the Seahorse XF HS Mini analyser before the assay run. Oligomycin (1.5 μM and Rotenone (0.5 μM) were loaded to the injection ports of the Seahorse XF real-time ATP rate assay sensor cartridge and replaced the utility plate with *Mtb* infected Seahorse XFp cell culture mini plate to start the assay. Wells with assay media alone were used as blank. Total 3 assay cycles were performed to record basal and mitochondrial inhibitors (Oligomycin and Rotenone) mediated oxygen consumption rate (OCR) and extra-cellular acidification rate (ECAR), which allows the calculation of total ATP production rate and pathway-specific dependency of cellular energy production. Assay was performed using the manufacturer’s user guide (Kit 103591-100).

### Bulk-RNA sequencing and Analysis

Total RNA was extracted from naïve and *Mtb* infected PKCδ^flox/flox^ and LysM^cre^PKCδ^flox/flox^ BMDM Qiagen RNeasy Mini Kit according to the manufacturer’s instructions. The integrity of RNA was assessed with Agilent Bioanalyzer DNA high sensitivity kit. RIN numbers were all above 8.4. RNA sequencing libraries were generated with KAPA RNA HyperPrep kit and sequenced with Illumina Nextseq2000 2x59-bp configuration targeting 30 million reads per sample. Sequencing reads were trimmed with fastp(*100*) to remove short reads and low quality 3’ end reads. Reads were aligned to mm10 genome with STAR(*101*) and low quality mappings, unmapped reads, multimappers were removed. Gene counts are obtained from filtered BAM files with HTSeq(*102*). Differential gene expression analysis was performed with DESeq2(*103*). Read counts were normalized for downstream analyses using the rlog transformation from DESeq2. An adjusted P value < 0.05 was used to determine significantly DEG. Volcano plots were generated with EnhancedVolcano package, while heatmap on “neutrophil migration” GO term genes was generated with pheatmap package. GO biological processes analysis was performed separately on upregulated or downregulated genes with |log2 fold change| > 0.5 and adjusted p value < 0.05 with ClusterProfiler package(*104*). Benjamini-Hochberg correction was applied to control False Discovery Rate and q-value 0.1 was used as a cutoff. RNA-Seq data are deposited in the National Center for Biotechnology Information (NCBI) Gene Expression Omnibus (GEO) and the reviewer token can be generated upon request.

### Liquid chromatography with tandem mass spectrometry (LC-MS/MS)

Mtb infected (MOI=1) and uninfected WT and LysM^cre^PKCδ^flox/flox^ BMDM cells (10 x 10^6^) were seeded and harvested in-plate using lysis buffer (1% SDS, 10mM tris-HCL pH 8, phosphatase and protease inhibitors Roche) and precipitated overnight at -20° C using 8:1:1 acetone: methanol: sample. Washed cell pellets were resuspended in denaturation buffer (6M urea, 2M thiourea, 10 mM Tris-HCL pH 8) and quantified using Bradford reagent. In-solution protein digestion was performed by reducing and alkylating 210 µg protein in 3 mM and 15 mM DDT and IAA for 20 minutes each, respectively. Urea is then diluted with 5 volumes of 10 mM Tris-HCL pH 8 followed by trypsin digestion (1:100) overnight at 30° C. digested peptides were then acidified to pH 2 and desalted by standard C18 SPE chromatography. Following elution, 10 µg peptides were aliquoted for proteome analysis with the remainder being used for phosphopeptide enrichment. Phosphopeptides were enriched using ReSyn Zirconium+ Magnetic Phosphopeptide Enrichment Beads (ReSyn). ReSyn Zirconium+ Magnetic Phosphopeptide Enrichment Beads were prepared by washing with 80% acetonitrile and 0.1% trifluoroacetic acid. The peptide samples were incubated with the prepared beads for 30 minutes at room temperature with gentle agitation. Subsequently, non-specifically bound peptides were removed by washing the beads three times with the same buffer. The phosphopeptides were then eluted with 1% ammonium hydroxide. Eluted phosphopeptides were acidified to pH 2 using 10% TFA and were desalted and concentrated before analysis by mass spectrometry. Proteomic and phospho-proteomic analysis was conducted using the Dionex Ultimate 3500 RSLC Nano System (Thermo Fisher) coupled to a Q Exactive mass spectrometer (ThermoFisher) as described previously(*105*). Raw files were processed and analysed using Perseus v1.6.5.0 (Maxquant) and MS/MS spectra were searched against the *Mus musculus* proteome database (http://www.uniprot.org/proteomes/UP000000589). Protein annotation (GO terms) and functional pathway enrichment (Reactome pathways) were conducted using Enrichr and GO Enrichment Analysis.

### Generation of human-monocyte-derived macrophages

Peripheral blood mononuclear cells (PBMC) were obtained from healthy donor buffy coats and were isolated using Lymphoprep (Alere Technologies) density gradient method. CD14 monocytes were isolated from PBMC through CD14^+^ magnetic bead separation kit (MACS Miltenyi) following the manufacturer’s protocol and differentiated at 1x10^6^ cells/ml in a 60 mm cell culture dish (Nunc, Denmark) for 7 days at 37°C in RPMI media (Sigma) supplemented with 1 mM Sodium Pyruvate (Sigma), 2 mM L-Glutamine (Sigma), 10% human AB Serum (hAB) (Sigma), and 50ng/ml recombinant human M-CSF (Biolegend). Following the 7-day differentiation, macrophages were then harvested by incubation with Accutase (Sigma) at 37°C for 20 minutes to gently detach the cells followed by centrifugation at 300 x g for 10 minutes. Cell pellets were resuspended in fresh prewarmed RPMI media (Sigma) supplemented with 1 mM Sodium Pyruvate (Sigma), 2 mM L-Glutamine (Sigma), 5% hAB (Sigma), counted, and seeded in desired tissue culture grade plates for further downstream experimental procedures.

### siRNA transfection

Human MDM were seeded at 80% confluency in a cell culture grade 96-well plate overnight at 37°C in a 5% CO2 incubator. TransIT-X2 transfection reagent (Mirusbio) and serum-free medium (Gibco) were prewarmed to room temperature. Next, Silencer-select siRNA (ThermoFisher) specific for PKCδ was added to the prewarmed serum-free medium followed by the addition of TransIT-X2 transfection reagent to allow the formation of the complex for 30 minutes at room temperature. The complex was then added dropwise to the seeded MDM at a final concentration of 25nM per well and incubated for 24-72 hours before continuing with *Mtb* infection. Knockdown efficiency of PKCδ was determined by qRT-PCR.

### Statistical analysis

All data were represented as mean values and analysed using GraphPad Prism v9.0. Statistical analyses were performed using an unpaired student t-test. Asterisks are defining significance compared to the control group as: *p < 0.05, **p < 0.01, ***p < 0.001, ****p < 0.0001.

## Acknowledgments

We thank Munadia Ansari for the maintenance and genotyping of mice. Faried Abbass for helping in aerosol *Mtb* infections. We are grateful to Lizzete Fick and Zoe Lotz for their excellent histology services.

## Author contributions

Conceptualization: RH, MO, SPP; Methodology: RH, MO, NP, SJ, SKLP, RR, SN, RMM, AFS, TG, A; Investigation: RH, MO, SPP; Visualization: RH, MO, TG, SPP; Supervision: RJW, FB, SPP; Writing original draft: RH, MO, SPP; Writing—review & editing: NDP, JB, MMM, RJW, FB, SPP.

## Data and materials availability

Further information and requests for resources and reagents should be directed to and will be fulfilled by the lead contact, Suraj Parihar. All data reported in this paper will be shared by the lead contact upon request. All original raw data have been deposited on GEO and are publicly available as of the publication date.

