## Supplementary Figures for "Protein Kinase C δ: a critical hub regulating macrophage immunomodulatory functions during *Mycobacterium tuberculosis* infection"

### Supplementary Fig 1.

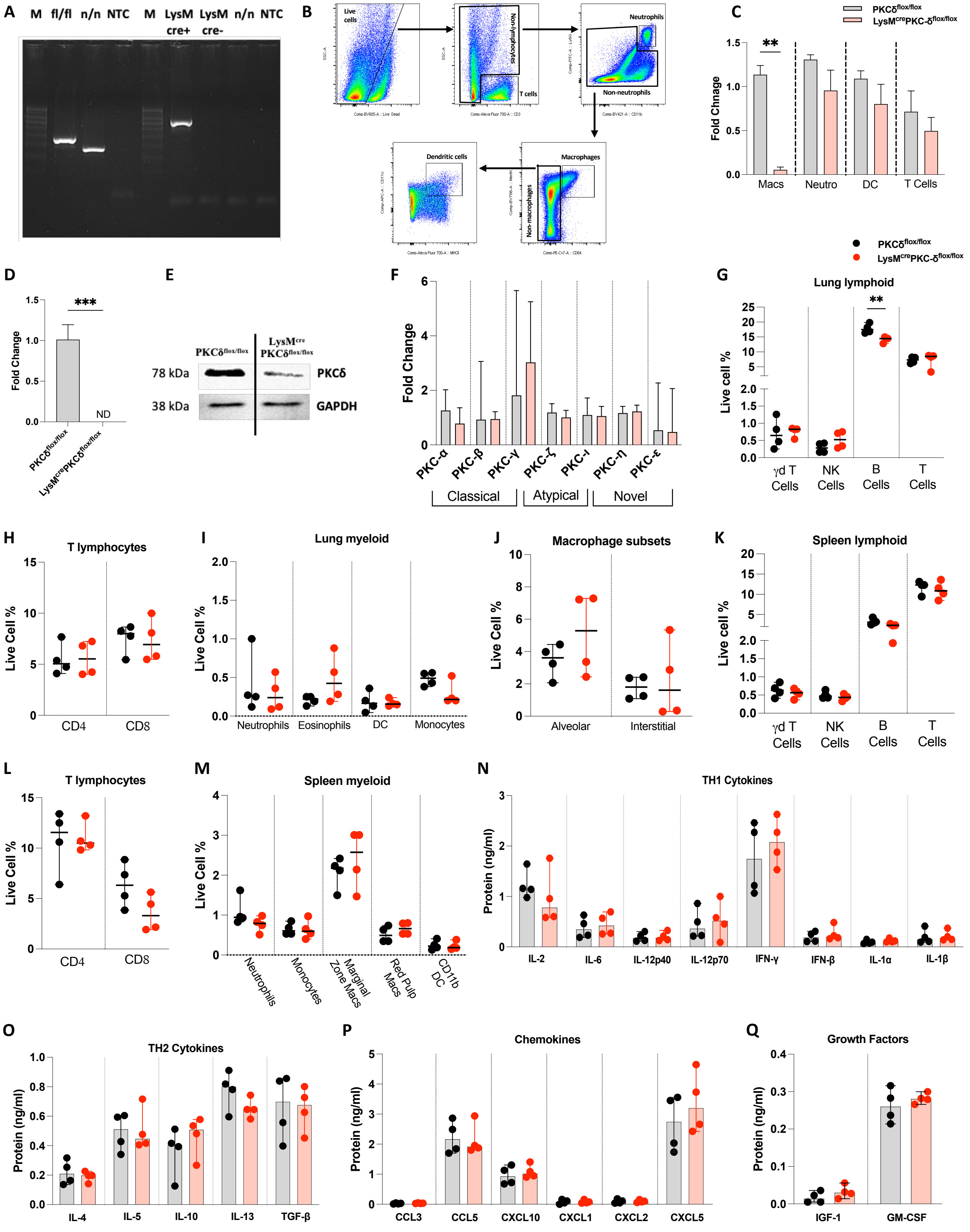

**Supplementary Figure 1. LysM$^{cre}$PKCδ$^{flox/flox}$ mice are indistinguishable from PKCδ$^{flox/flox}$ littermate control at the naive state. [A]** Integrity of loxP-floxed PKCδ and presence of LysMcre was confirmed in the genomic DNA extracted from the tail cuts of PKCδ$^{flox/flox}$ and LysM$^{cre}$PKCδ$^{flox/flox}$ mice. Representative agarose gel demonstrates the marker (m), wild-type (n/n), floxed PKCδ (fl/fl), LysM$^{cre}$PKCδ$^{flox/flox}$ (LysMcre+), PKCδ$^{flox/flox}$ (LysMcre-), non-template control (NTC). **[B]** Gating strategy for lung sorted immune cells and confirmation of PKCδ deletion in macrophages by qRT-PCR. **[C]** PKCδ expression was determined in sorted immune cell populations by qRT-PCR. Expression levels are normalized to the endogenous housekeeping gene Hprt. **[D-E]** Deletion of PKCδ in bone- marrow derived macrophages was confirmed by qRT-PCR and western blot analysis. HPRT and GAPDH were used as a normalizing control in qRT-PCR and Western blot respectively. **[F]** Protein kinase C isoform-specific primers were used for gene expression analysis in PKCδ deficient bone-marrow derived macrophages (LysM$^{cre}$PKCδ$^{flox/flox}$) as compared to the control (PKCδ$^{flox/flox}$) by qRT-PCR. **[G-M]** Various lymphoid and myeloid immune populations were detected by flow cytometry in the lung [G-J] and spleen [K-M] in both PKCδ$^{flox/flox}$ and LysM$^{cre}$PKCδ$^{flox/flox}$ mice. **[N-Q]** Determining cytokine levels in lung homogenates collected from PKCδ$^{flox/flox}$ and LysM$^{cre}$PKCδ$^{flox/flox}$ mice. All data shown mean+/- SD and is representative of two independent experiments with n=4 mice/group. Statistical analyses were performed using an unpaired student t-test. Asterisks are defining significance compared to the control group as: **$p < 0.01$, ***$p < 0.001$.

Supplementary Fig 2.

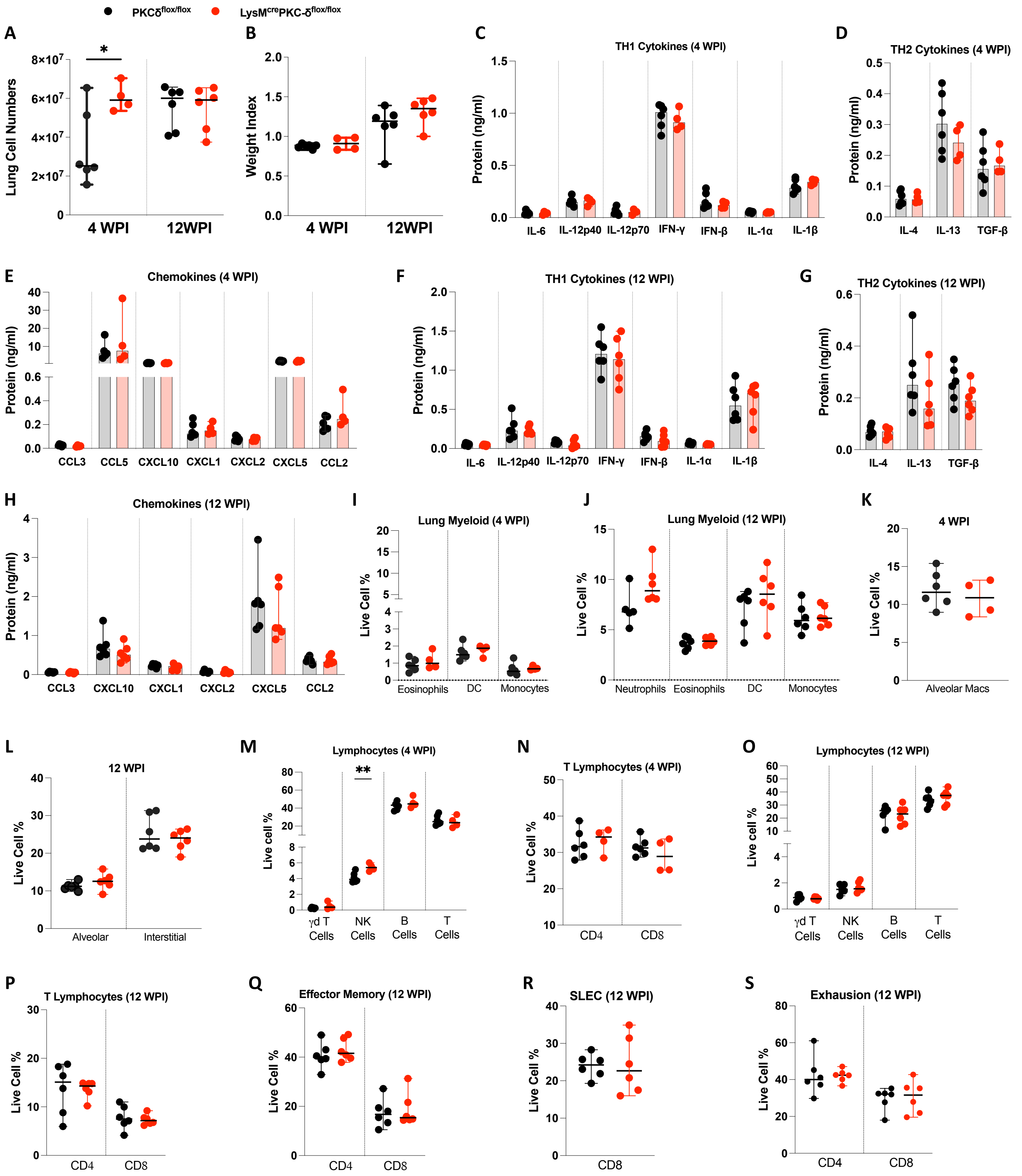

**Supplementary Figure 2. Lung cell numbers, cytokine profiling, and immune cell phenotyping of LysM$^{cre}$PKCδ$^{flox/flox}$ mice during acute (4WPI) and chronic (12 WPI) *Mtb* infection. [A]** Lung cell numbers and **[B]** weight index were determined at acute and chronic stage of *Mtb* infection. **[C-H]** Determining cytokine and chemokine levels in lung homogenates collected from PKCδ$^{flox/flox}$ and LysM$^{cre}$PKCδ$^{flox/flox}$ mice at acute and chronic stage of *Mtb* infection. **[I-S]** Various myeloid and lymphoid immune cell populations were detected by flow cytometry at acute and chronic stage of *Mtb* infection. All data shown mean+/- SD and is representative of two independent experiments with n=4-6 mice/group. Statistical analyses were performed using an unpaired student t-test. Asterisks are defining significance compared to the control group as: *$p < 0.05$, **$p < 0.01$.

**Supplementary Fig 3.**

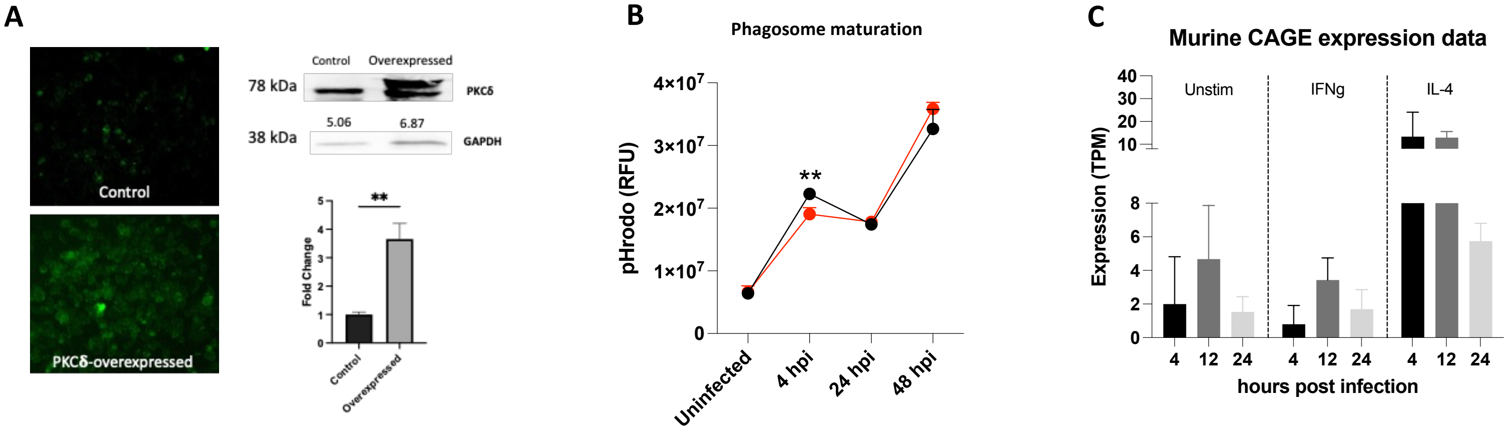


**Supplementary Figure.3. PKCδ overexpression in RAW264.7 murine macrophages, determination of phagosome maturation in LysM$^{cre}$PKCδ$^{flox/flox}$ BMDMs and murine PKCδ expression during *Mtb* infection (adapted from FANTOM5 CAGE database). [A]** Lentivirus-mediated overexpression of PKCδ in RAW264.7 murine macrophage cell line confirmed by ZOE fluorescence imaging, western blot, and qRT-PCR. **[B]** Determination of phagosome maturation (Relative fluorescence unit) based on pH change in the phagolysosomal compartment in bone-marrow-derived macrophages from LysM$^{cre}$PKCδ$^{flox/flox}$ and PKCδ$^{flox/flox}$ mice. **[C]** Determining PKCδ expression in classically (IFNg-stimulated) or alternatively (IL-4 stimulated) activated murine macrophages utilizing publicly available FANTOM5 CAGE database. Asterisks are defining significance compared to the control group as: $^{**}p < 0.01$.

**Supplementary Fig 4.**

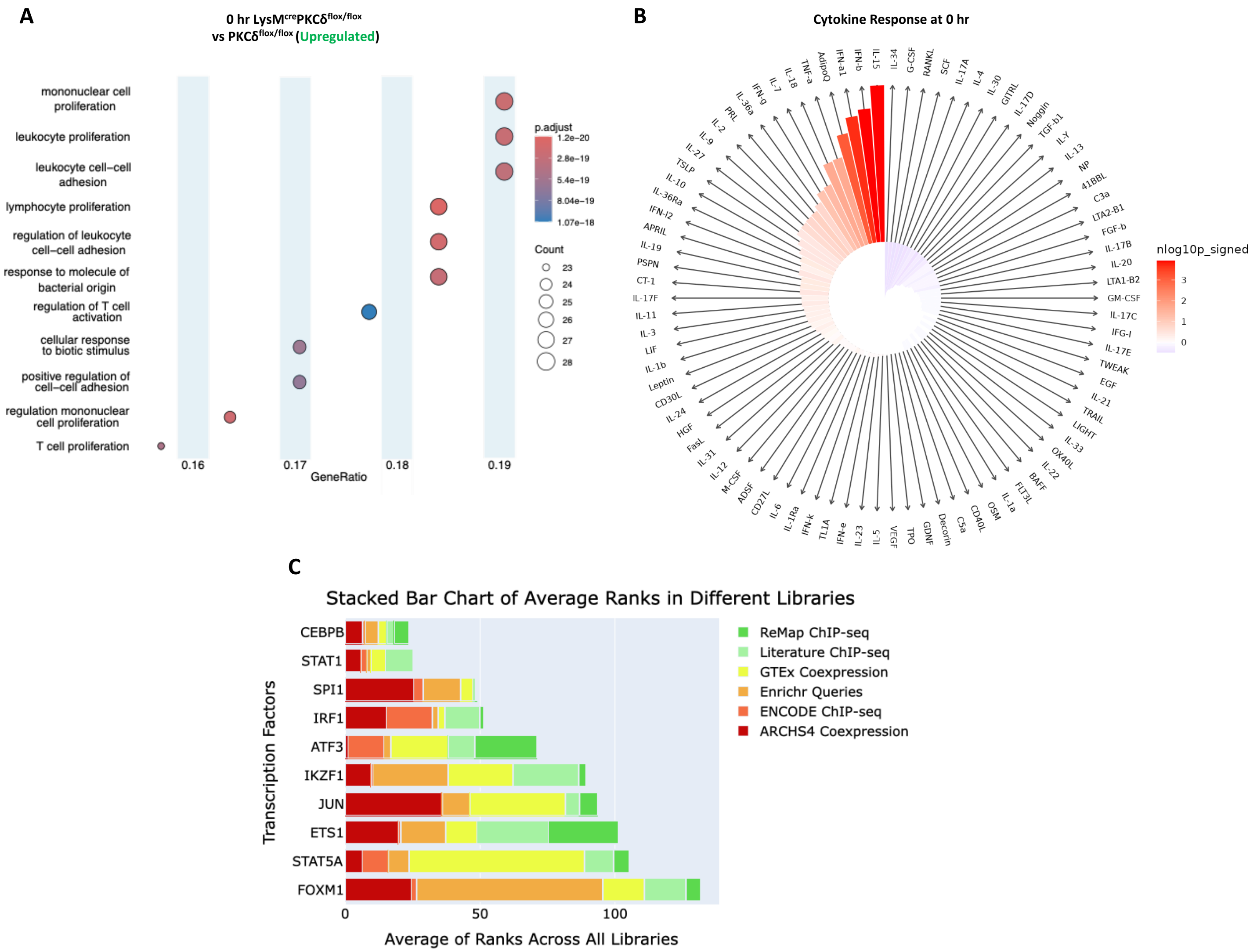

**Supplementary Figure.4. Gene ontology, cytokine response (IREA), and ChEA analysis in the LysM$^{cre}$PKCδ$^{flox/flox}$ BMDMs at naive state. [A]** Horizontal dot plots (Ora) detailing the association of enriched Gene Ontology (GO) biological processes are shown between LysM$^{cre}$PKCδ$^{flox/flox}$ and PKCδ$^{flox/flox}$ BMDMs at 0 hr (uninfected) time point. **[B]** IREA cytokine enrichment plot showing the enrichment score (ES) for each of the cytokine response in LysM$^{cre}$PKCδ$^{flox/flox}$ BMDMs at 0 hr (uninfected) time point. Bar length is representing the ES with darker red (enriched in LysM$^{cre}$PKCδ$^{flox/flox}$ BMDMs) and darker blue (enriched in PKCδ$^{flox/flox}$ BMDMs). **[C]** Horizontal bar chart representing the top ranked transcription factors at 0 hr (uninfected) time point according to their average integrated scores across all the libraries. All data shown are analysed and produced using R studio packages and appyters web-based software.

**Supplementary Fig 5.**

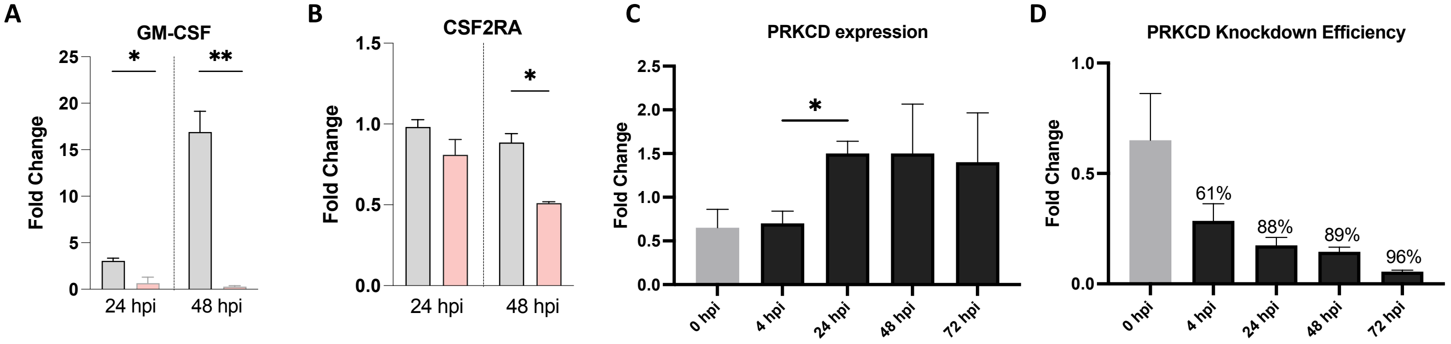


**Supplementary Figure.5. mRNA expression of GM-CSF and CSF2RA in LysM$^{cre}$PKCδ$^{flox/flox}$ BMDMs, mRNA expression of PKCδ in MDMs and siRNA mediated PKCδ knockdown efficiency in MDMs during *Mtb* infection. [A-B]** mRNA expression of GM-CSF and CSF2RA in bone-marrow-derived macrophages from LysM$^{cre}$PKCδ$^{flox/flox}$ and PKCδ$^{flox/flox}$ mice during *Mtb* infection at indicated time points. **[C]** mRNA expression of PRKCD in human monocyte-derived macrophages at indicated time points during *Mtb* infection. **[D]** siRNA mediated knockdown efficiency of PRKCD in human monocyte-derived macrophages during *Mtb* infection. Asterisks are defining significance compared to the control group as: *p < 0.05, **p < 0.01.

**Table S1.**

| Gene | Accession number | Forward Primer (5'-3') | Reverse Primer (5'-3') |
|---|---|---|---|
| **Mouse PKCδ** | M69042 | CTGGGTAACTTAACAAGACC | CTGCTAAATAACATGTTCGGTCC |
| **Mouse PKCε** | M18331 | CATCGATCTCTCGGGATCATCG | CGGTTGTCAAATGACAAGGCC |
| **Mouse PKCη** | M62980 | AGCTAGCCGTCTTCCACGAGACGC | GGACGACGCAGGTGCACACTTGG |
| **Mouse PKCθ** | D11061 | AGCTAGCCGTCTTCCACGAGACGC | GGACGACGCAGGTGCACACTTGG |
| **Mouse caspase 1** | NM_009807 | ACAAGGCACGGGACCTATG | TCCCAGTCAGTCCTGGAAATG |
| **Mouse caspase 11** | NM_009807 | AGAGGGCATGGAGTCAGAGA | GCCATGAGACATTAGCACCA |
| **Mouse NLRP3** | NM_145827 | ATTACCCGCCCGAGAAAGG | TCGCAGCAAAGATCCACACAG |
| **Mouse AIM2** | NM_001013779 | GTCACCAGTTCCTCAGTTGTG | CACCTCCATTGTCCCTGTTTTAT |
| **Mouse IL-1β** | NM_008361 | GCTTCAGGCAGGCAGTATC | AGGATGGGCTCTTCTTCAAAG |
| **Mouse IL-18** | NM_008360 | ACTTTGGCCGACTTCACTGT | GGGTTCACTGGCACTTTGAT |
| **Mouse GM-CSF** | XM_006532127 | ACCACCTATGCGGATTTCAT | TCATTACGCAGGCACAAAAC |
| **Mouse CSF2RA** | NM_009970 | ACGTGGCGCGATGCAT | ACTTGTCACTGCTGGGGAGTG |
| **Mouse iNOS** | NM_011198 | AGCCCTCACCTACTTCCTG | CAATCTCTGCCTATCCGTCTC |
| **Human PRKCD** | L07860 | CACCATCTTCCAGAAAGAACG | CTTGCCATAGGTCCCGTTGTTG |
